# The dynamic interplay between tonic and phasic arousal shapes attention to optimize performance

**DOI:** 10.1101/2024.03.22.586259

**Authors:** Aurélie Grandjean, Anne Mathieu, Lilia Ponselle, Sophie Chen, Andreas Widmann, Nicole Wetzel, Aurélie Bidet-Caulet

## Abstract

Arousal and attention are fundamental brain functions that play a critical role in optimizing performance. Kahneman’s attention model (1973) theorizes a key interplay between attention and arousal, yet this relationship remains poorly understood. Using a multimodal approach, we investigated this interaction in healthy young adults performing an auditory attention task designed to simultaneously assess phasic arousal, voluntary attention, and involuntary attention. Furthermore, tonic arousal was experimentally modulated with low or high arousing music, as confirmed by changes in pupil size and heart rate. Behavioral data confirmed that informative cues enhanced voluntary attention, while unexpected salient task-irrelevant sounds (so-called distractors) produced either shortened or lengthened reaction times depending on their timing relative to target onset. Physiological data indicated that the facilitation effect of distractors on reaction times was driven by increases in phasic arousal. This benefit was further modulated by a dynamic interplay between phasic arousal and voluntary attention over time. Supporting Aston-Jones and Cohen’s theory, a fronto-central cortical response to distractors revealed that tonic and phasic arousal interact in line with an inverted U-shaped relationship. This study provides empirical evidence, in humans, that tonic arousal can optimize performance by tuning phasic arousal and attentional control.

## 1. INTRODUCTION

Crucial for the selection and prioritization of sensory information, attention enables individuals to navigate their complex surroundings efficiently. It is a multifaceted cognitive process, manifesting in two primary forms: voluntary and involuntary attention. The former facilitates focused engagement with specific tasks, encompassing processes such as voluntary orienting, anticipation, and sustained attention. In contrast, involuntary attention is elicited by salient, unexpected stimuli outside the intended focus. The intricate balance between these two aspects is essential for an optimal attentional control. Over the last 50 years, several theoretical and physiological models of attention have been proposed (Broadbent, 1971; Corbetta et al., 2008; Corbetta & Shulman, 2002; Kahneman, 1973; Mesulam, 1981; Posner & Petersen, 1990), many of them emphasizing the interplay between attention and arousal. Arousal is generally described as the state of physiological reactivity of a subject (Broadbent, 1971; Coull, 1998; Kahneman, 1973; Näätänen, 1992; Sturm & Willmes, 2001), and more recently as a general state of excitability of the cortex, modulated by the activity of the Locus Coeruleus – Norepinephrine (LC-NE) system (Aston-Jones & Cohen, 2005; Nieuwenhuis et al., 2011).

Since Yerkes and Dodson (1908) pioneer works in mice, an inverted U-shaped relationship between arousal and performance has been proposed, with moderate levels of arousal promoting maximal performance, while lower and upper levels would lead to poorer performance. This inverted U-shaped relationship was later associated to visual voluntary attention by Easterbrook (1959) in a review of behavioral findings: at a moderate level of arousal, attention would be focused on relevant information while irrelevant one could be ignored. In hypo- or hyper-arousal states, relevant information would become indistinguishable from irrelevant information. Importantly, the optimal arousal level would be higher for simple than for complex tasks. In 1973, Kahneman (Kahneman, 1973) proposed a comprehensive capacity model of attention encompassing both transient (i.e., phasic) and sustained (i.e., tonic) arousal. This model suggests that available attentional capacities and arousal can influence each other to achieve optimal performance. This model also considers that the orienting response to novel stimuli encompasses both involuntary allocation of attention to the novel and a transient increase in arousal (see also: Näätänen, 1992, for a similar proposal). Later on, based on neuropsychological observations, Mesulam (1981) proposed a strong influence of the reticular system underlying arousal on both sensory and higher-level brain regions in the frontal and parietal cortices involved in attention, a model regularly refined over the following decades (Corbetta et al., 2008; Corbetta & Shulman, 2002; Posner & Petersen, 1990).

These hypotheses were confirmed by experimental evidence, mostly in rodents and monkeys, showing widespread projections from the LC-NE system to the cerebral cortex, the cerebellum, and subcortical structures (thalamus, amygdala) (Berridge & Waterhouse, 2003). Conversely, attentional control has been shown to be supported by brain networks including frontal and parietal regions (Corbetta & Shulman, 2002; Posner & Petersen, 1990), with descending connections to the locus coeruleus (Corbetta et al., 2008). Animal studies have demonstrated that neurons in the LC present two different modes of activity: a tonic mode, that defines the baseline activity of the system and changes relatively slowly, and a phasic mode during which neurons fire at a high frequency (10–15 spikes per second) in response to a salient stimulus (Vazey et al., 2018). Tonic firing of LC neurons is relatively low when the organism is at rest, medium when engaged in a task requiring the filtering of irrelevant information, and high when the exploration of the environment requires to stay attentive to potentially unexpected occurring events (for a review see: Aston-Jones & Cohen, 2005). Therefore, experimental findings confirm that the LC-NE system presents both phasic and tonic activities which can modulate or be modulated by signal from cortical regions supporting attention. However, the interplay between attention and arousal remains poorly understood, in particular in humans, and is still at the heart of recent influential theories of task performance and stimuli processing (Aston-Jones & Cohen, 2005; Berridge & Waterhouse, 2003; Mather et al., 2016).

Involuntary allocation of attention to novel stimuli can be behaviorally measured by a cost in performance (longer reaction times and reduced accuracy), named distraction effect (Wetzel & Schröger, 2014). However, an increasing number of studies have reported improved performance when novel sounds are presented (SanMiguel et al., 2010; Wetzel et al., 2012), particularly in the context of highly arousing emotional stimuli (Max et al., 2015). This facilitation effect has been attributed to an increase in phasic arousal triggered by the distracting sound (Bidet-Caulet et al., 2015; Masson & Bidet-Caulet, 2019), in line with attention models of the orienting response (Kahneman, 1973; Näätänen, 1992) and animal studies showing an increase in LC-NE activity in response to salient stimuli (Aston-Jones et al., 1991; Devilbiss, 2019; Grant et al., 1988; Rasmussen et al., 1986). At the cerebral level, novel unexpected sounds elicit a distinctive sequence of event-related potentials (ERPs): the N1/Mismatch Negativity around 100ms, followed by a P3 complex around 300ms (also called novelty-P3 or P3a) (Escera et al., 1998; Näätänen, 1992). The P3 complex is commonly thought to reflect the involuntary orienting of attention towards distracting stimuli (Escera et al., 1998; Polich & Criado, 2006). Moreover, this P3 complex is dissociable in two phases: a fronto-central early phase peaking around 235ms called early-P3 and a fronto-parietal late phase peaking around 320ms called late-P3 (Yago et al., 2003). Recently, the amplitude of the early-P3 was found strongly correlated to the arousal rating of distracting sounds (Masson & Bidet-Caulet, 2019; Widmann et al., 2018), suggesting that the early-P3 would reflect the arousal component of the orienting response. In the last few years, an increasing number of studies used pupillometry recordings to investigate attentional processes and showed that novel oddball sounds cause a transient dilation of the pupil higher for arousing emotional stimuli, presumably due to the co-activation of the sympathetic nervous and the LC-NE systems (Widmann et al., 2018).

Several experimental studies have investigated the link between tonic arousal and voluntary attention. The influential Aston-Jones theory of the LC-NE function in cognition relies mainly on animal studies. They were able to record LC neuron activity using electrophysiology in monkeys, and correlate levels of tonic and phasic activity with performance during visual attention tasks (Aston-Jones et al., 1991, 1999; Usher et al., 1999). They confirmed the Yerkes and Dodson inverted U-shaped relationship with moderate levels of tonic arousal promoting strong phasic activity in the LC-NE and maximal performance; and lower and upper levels resulting in small or no phasic LC-NE activity and poorer performance. In the human literature, for technical reasons, evidence is much rarer and mainly in the visual modality. Tonic arousal modulations are achieved using a wide range of methodologies: pharmacological manipulations (Coull et al., 2004), stress induction (Rojas-Thomas et al., 2023), meditation (Kozhevnikov et al., 2022), isometric exercise (Mather et al., 2020) or background music presentation (Cloutier et al., 2020; Kiss & Linnell, 2021; Nadon et al., 2021). In addition, some behavioral studies have used pre-task music to induce tonic arousal modulation, with varying type and duration of music extracts (Dovorany et al., 2023; Quan et al., 2023). Because arousal level in human cannot be assessed directly by measuring the activity of neurons within the LC, most studies have relied on subjective ratings of the subject arousal state (McConnell & Shore, 2011) or on indirect physiological markers such as heart rate or salivary cortisol levels (Rojas-Thomas et al., 2023). The obtained behavioral results are quite mixed, with for example relaxing (i.e. low arousing) music being beneficial to the performance of a Stroop test (Nadon et al., 2021); while impairing performance in a visuo-spatial flanker task (Cloutier et al., 2020). Very few studies have investigated the link between tonic arousal, phasic arousal and attention at the brain level (Mather et al., 2020; Rojas-Thomas et al., 2023). Interestingly, Mather et al. (Mather et al., 2020) conducted a fMRI study in which isometric exercises were used to modulate tonic arousal shortly before an auditory oddball task. Handgrip was found to decrease tonic pupil size during the subsequent task while increasing phasic pupil responses to oddball sounds, increasing activation of the frontoparietal network and speeding oddball detection, suggesting that reducing tonic arousal increases phasic arousal and activation of the brain network of voluntary attention.

Therefore, the interplay between arousal and attention remains largely unexplored, in particular in humans. To our knowledge, the effect of tonic or phasic arousal levels on voluntary and involuntary attention has not been directly addressed. In this study, we propose to unravel the complex relationship between voluntary attention, involuntary attention, tonic and phasic arousal. For this purpose, we used the Competitive Attention Task (CAT) which provides measures of voluntary and involuntary attention processes and phasic arousal (Bidet-Caulet et al., 2015). In the CAT, voluntary attention orienting is modulated by modifying the information provided by a cue preceding a target sound to detect. Involuntary attention and phasic arousal are triggered by rare, unexpected task-irrelevant sounds (distracting sounds) presented at random times between the cue and the target (**Fig. 1**). Phasic arousal is further modulated by varying the arousing content of the distracting sounds. Different levels of tonic arousal are induced by presenting calm or dynamic short music tracks to the participants before each block of the CAT. The level of tonic arousal is indirectly assessed using peripheral measures of autonomic nervous system activity such as the heart rate, skin conductance and pupil size (Bellato et al., 2022; Wass et al., 2015). The pupil dilation response (PDR) to the different events of the CAT is used as an indirect measure of phasic arousal (Gilzenrat et al., 2010; Joshi et al., 2016; Nieuwenhuis et al., 2011), while EEG measures are used to assess brain processes involved in voluntary and involuntary attention and phasic arousal (Bidet-Caulet et al., 2015). According to theoretical models of attention and previous experimental findings, we hypothesize that a moderate level of tonic arousal induced with low arousing music – compared to a higher level – will facilitate attention effects, consistent with Kahneman’s capacity model of attention (Kahneman, 1973). We further expect this moderate tonic arousal to increase the phasic arousal response to distracting sounds, in line with the adaptive gain and optimal performance theory proposed by Aston-Jones and Cohen (Aston-Jones & Cohen, 2005). We therefore anticipate that a moderate level of tonic arousal would result in optimal performance by tuning phasic arousal and attentional control. Additionally, we hypothesize that distracting sounds will elicit an increase in phasic arousal, which will be further amplified by highly arousing content, leading to improved behavioral performance.

**Fig. 1.**
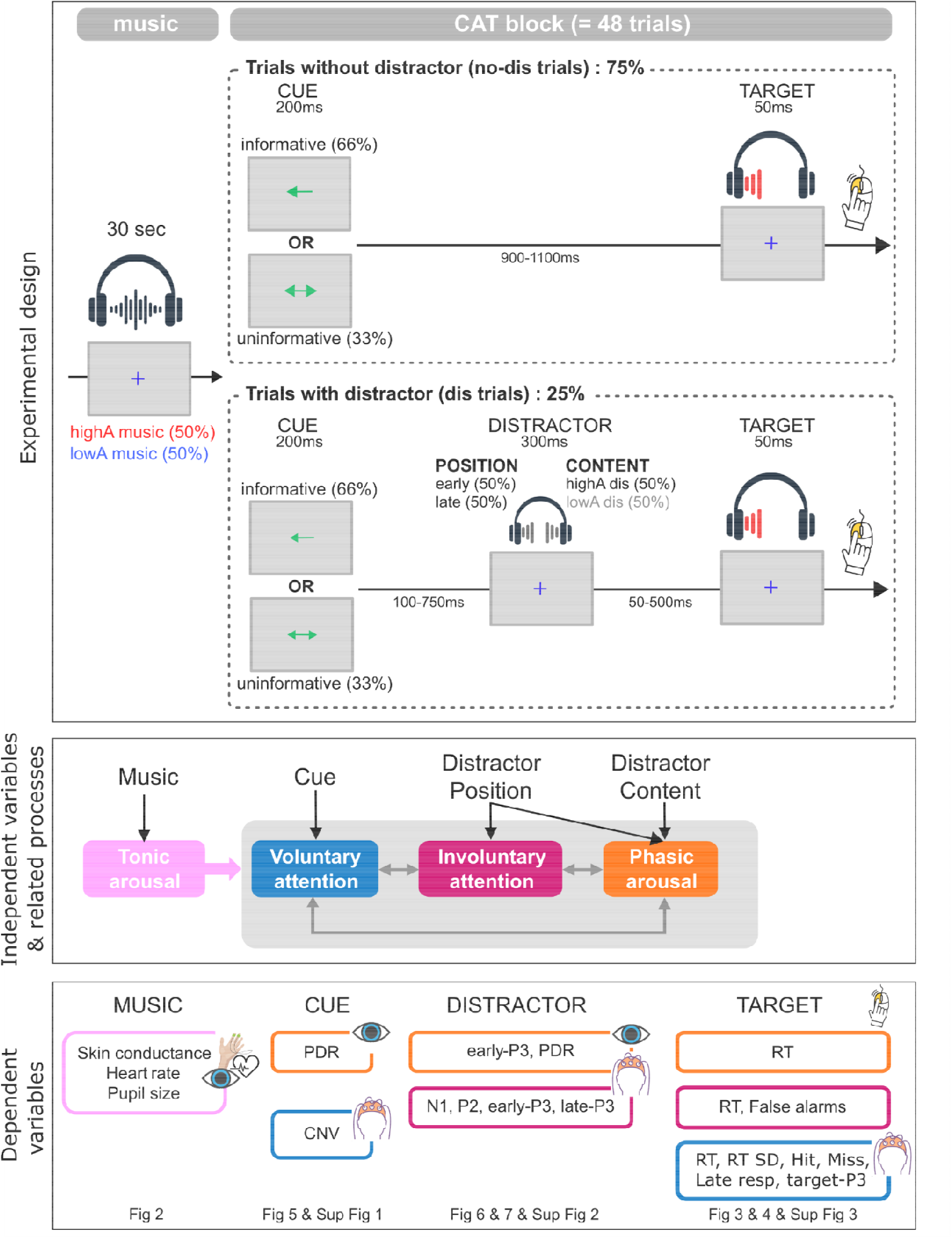
Protocol. Experimental design. Each block of the Competitive Attention Task (CAT; 48 trials) is preceded by 30 sec of music (**high arousing: highA music**, or **low arousing: lowA music**). In trials without distractor (no dis, top), a cue (**informative** or **uninformative**) is first presented on the screen, and after a delay (between 900 and 1100ms) a target sound is played in the left or right ear. In trials with distractor (bottom), a distractor sound is presented binaurally in the delay between the cue and the target sound. Distractors presented between 600 and 800ms before target onset cue are called **early distractors** (**early dis**). Distractors presented between 350 and 550ms before target onset are called **late distractors** (**late dis**). These distracting sounds can be either **high arousing** (**highA dis**), or **low arousing** (*lowA dis*). Subjects were asked to press the mouse button as fast as possible when they hear the target sound. **Independent variables and related brain processes**. High and low arousing **music** were used to modulate the level of **tonic arousal**. Informative compared to uninformative **cues** increase *voluntary attention* to the task. **Distractors** trigger both *involuntary attention* and an increase in *phasic arousal* that can be dissociated according to the **distractor position**. The **arousing distractor content** allows further modulation of *phasic arousal*. This design allows investigation of the interaction between involuntary attention, voluntary attention and phasic arousal, as well as the impact of tonic arousal on these interactions. **Dependent variables**. The impact of the independent variables was investigated on different physiological or behavioral measures locked to the different events of the paradigm and associated with distinct brain processes (see color correspondence with panel b). CNV: contingent negative variation, PDR: pupil dilation response, RT: reaction time, RT SD: reaction time standard deviation.

## 2. RESULTS

The impact of the tonic arousal level, induced by presenting low arousing (lowA) or high arousing (highA) music before each block of the Competitive Attention Task (CAT, Fig. 1), was explored on peripheral measures (skin conductance, pupil size and heart rate), as well as on performance, event-related pupil dilation response (PDR) and event-related potential (ERP) during the CAT. We used Bayesian Wilcoxon tests when only the effect of music was tested. We used linear mixed models (LMM) to estimate the effect of the independent variables (Music, Cue, Distractor Position, Distractor Content). The model characteristics are presented in Table 1.

**Table 1.**
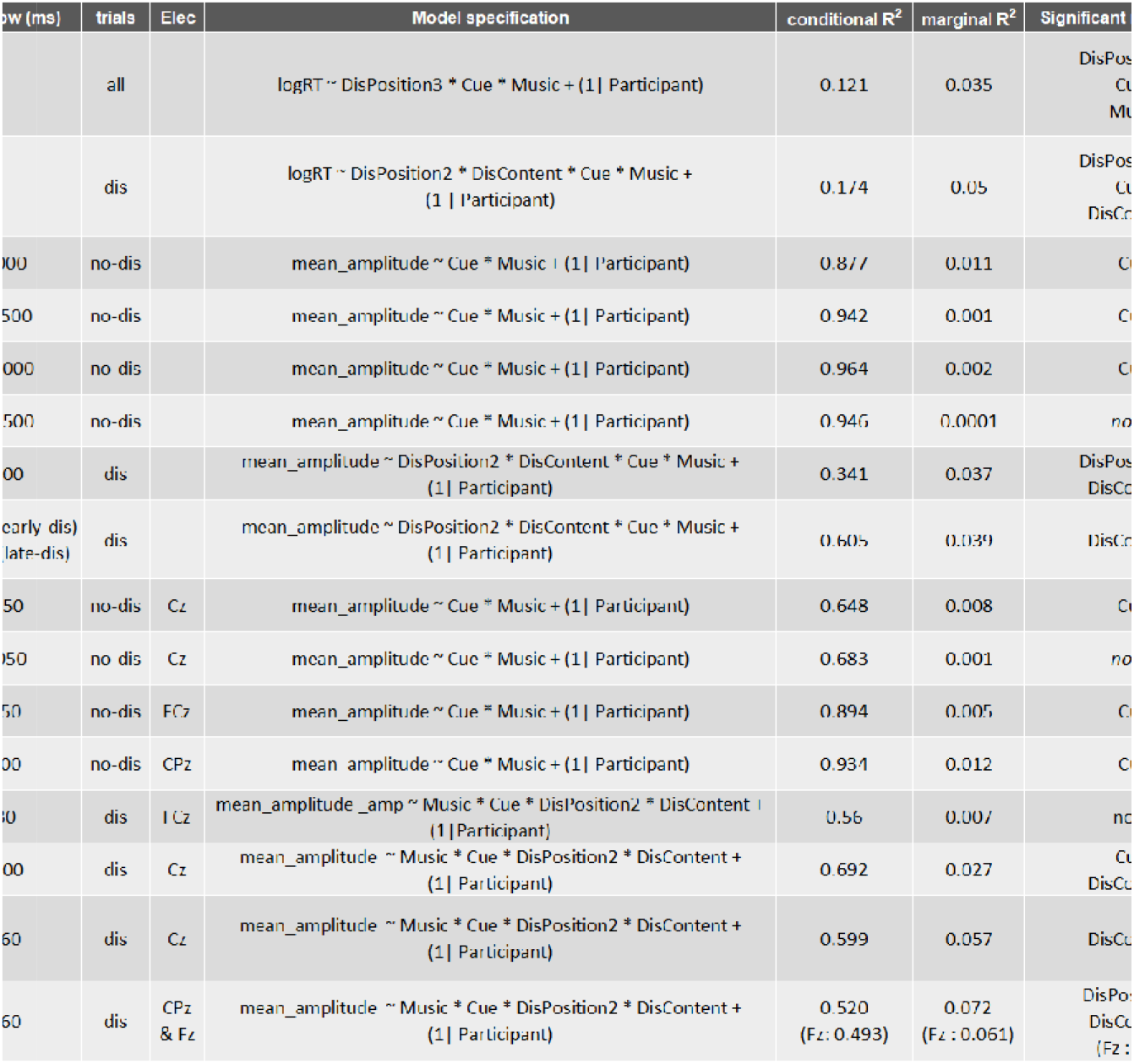
Details of the statistical models for each analysis. For each dependent variable, are provided details of the mixed model (fixed within-subject factors and random factors), information on model performance (marginal and conditional R^2^ when available) and results of the statistical analyses (significant main effect and significant interactions). Factors levels: Music (2 levels: highA music, lowA music), Cue (2 levels: informative, uninformative), DisPosition2 (2 levels: early, late), DisPosition3 (3 levels: no dis, early, late), and DisContent (2 levels: highA dis, lowA dis). RT+: positive RT (RT > 0ms).

### 2.1 Arousing music enhances peripheral measures of tonic arousal

To validate the modulation of the tonic arousal level, we explored the effect of the music type on peripheral measures (Fig. 2). We didn’t find evidence for an effect of music on skin conductance (BF_10_ = 1.09), but we found strong to decisive evidence for an effect of music on pupil size (BF_10_ = 33.87) and heart rate (BF_10_ = 520.23). These two measures were larger during the listening of high arousing (highA) music compared to low arousing (lowA) music (mean ± SEM: change of pupil size, highA: −0.01 ± 0.02 mm & lowA: −0.08 ± 0.02 mm; heart rate, highA: 70.0 ± 0.5 bpm & lowA: 69.2 ± 0.5 bpm). Two out of three physiological measures are enhanced during listening to high arousing music compared to low arousing music, validating the tonic arousal modulation.

**Fig. 2.**
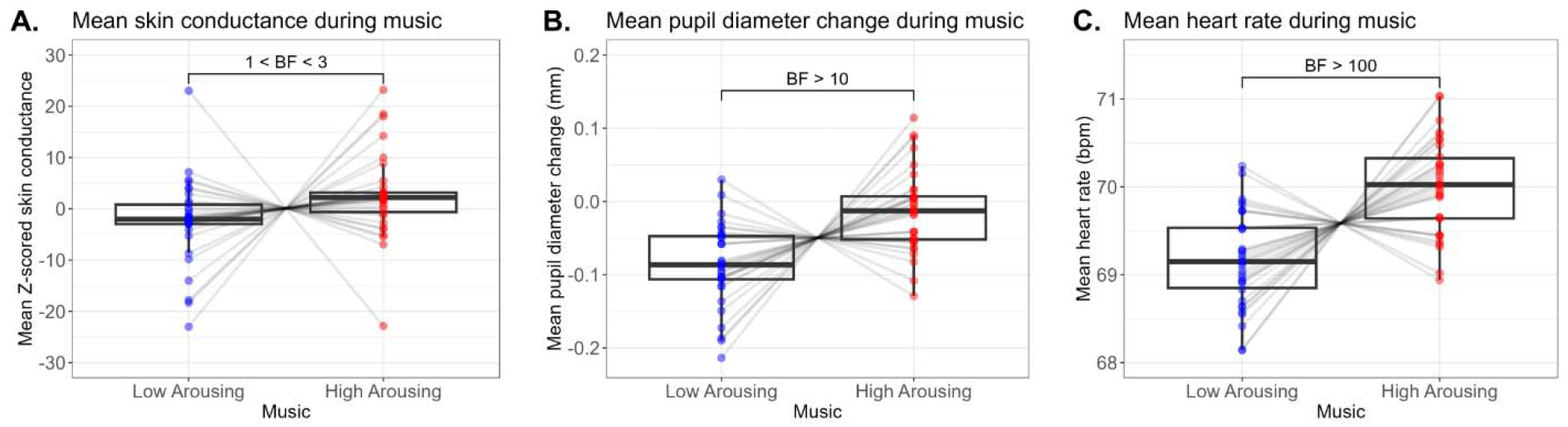
**Physiological measures during music listening**, as a function of music type. **A**. Mean Z-scored skin conductance. **B**. Mean pupil size change from baseline in millimeters (mm). **C**. Mean heart rate in beat per minute (bpm). Boxplots were plotted using normalized individual data (= raw data - subject mean + group mean). Within each boxplot, the horizontal line represents the group median, the lower and upper hinges correspond to the first and third quartiles. The upper whisker extends from the hinge to the largest value no further than 1.5 * IQR from the hinge (IQR = inter-quartile range, or distance between the first and third quartiles). The lower whisker extends from the hinge to the smallest value at most 1.5 * IQR of the hinge. Superimposed to each boxplot, the dots represent individual means. BF: Bayes Factor.

### 2.2. Music impacts behavioral measures of attention and phasic arousal

#### 2.2.1. Reaction Times are shorter after low arousing music and increased phasic arousal

Reaction times (RT) were fitted to a linear mixed model with Music, Cue and DisPosition3 (no, early or late distractor). Consistent with previous works (Bidet-Caulet et al., 2015; Hoyer et al., 2021; Hoyer, Abdoun, et al., 2023), RT were significantly modulated by the type of Cue (χ^2^ (1) = 316.93, p < 0.001, ω^2^⍰: very small) and the position of the Distractor (χ^2^ (2) = 1174.34, p < 0.001, ω^2^⍰: small). RT were shorter in the informative cue condition (mean ± SEM: 255.2 ± 0.7 ms) compared to the uninformative cue condition (273.6 ± 1.0 ms). RT were shorter in the early distractor (dis) condition (226.1 ± 1.4 ms) compared to the late dis condition (268.5 ± 1.8 ms) and to the no dis condition (265.8 ± 0.6 ms).

RT were also significantly modulated by the type of Music (χ^2^ (1) = 6.05, p < 0.05, ω^2^⍰: very small). RT were shorter in the low arousing music condition (261.0 ± 0.8 ms) compared to the high arousing music condition (261.8 ms ± 0.8 ms).

Importantly, significant interactions were also revealed: Cue x Music (χ^2^ (1) = 4.30, p < 0.05, ω^2^1]: very small) and DisPosition3 x Cue x Music (χ^2^ (2) = 7.23, p < 0.05, ω^2^1]: very small). Post-Hoc HSD tests were conducted to analyze this triple interaction (Fig. 3). In the high arousing music condition, the cue effect was significant in the no dis (p < 0.0001, d: small), early (p < 0.0001, d: small) and late (p < 0.001, d: very small) dis conditions; while in the low arousing music condition, the cue effect was only significant in the no (p < 0.0001, d: very small) and late (p < 0.001, d: very small) dis conditions. Moreover, in trials with early distractor, the RT were shorter after low arousing music than after high arousing music after uninformative cues, only.

**Fig. 3.**
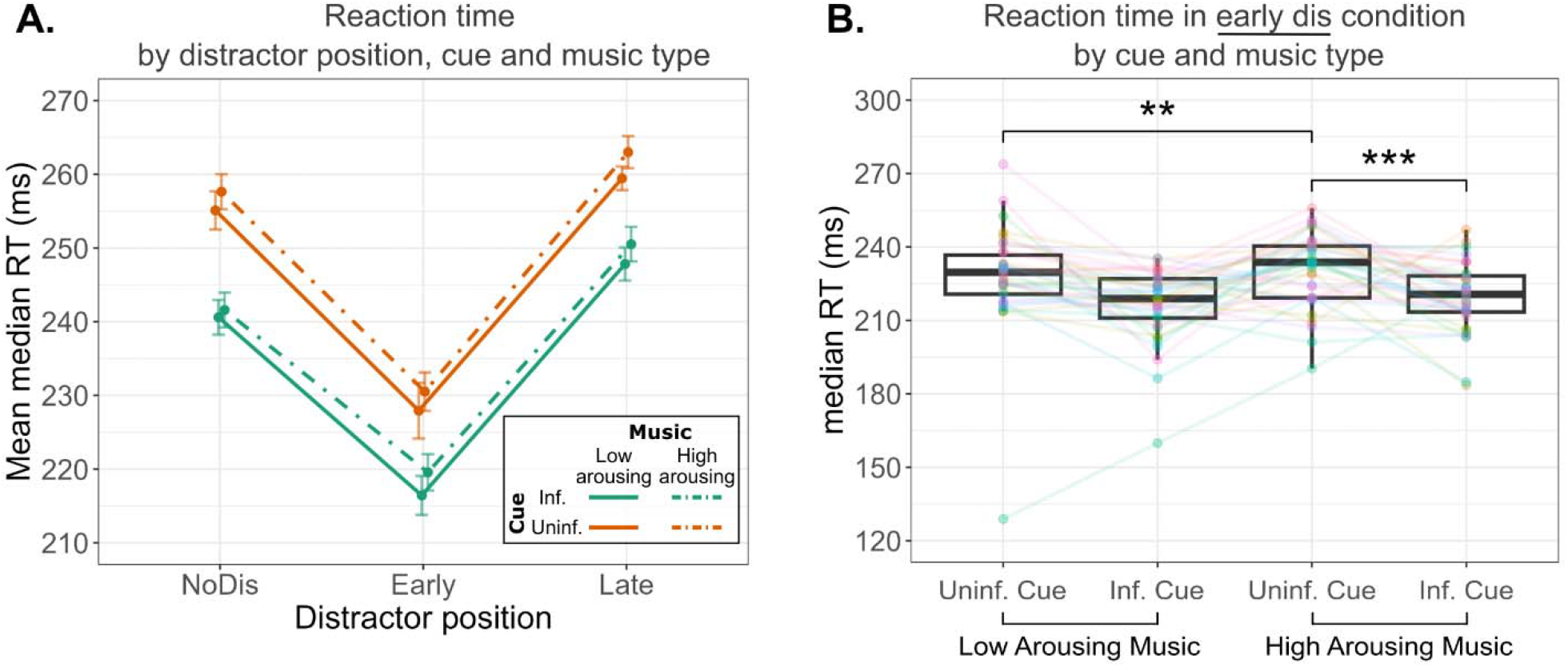
RT results. **A**. Mean RT by distractor position, cue and music type. Each point corresponds to values averaged across subjects. Within-subject standard errors of the mean are plotted around group means. **B**. Median RT by cue and music type, in trials with early distractors. Boxplots were plotted using normalized individual data (= raw data - subject mean + group mean). Within each boxplot, the horizontal line represents the group median, the lower and upper hinges correspond to the first and third quartiles. The upper whisker extends from the hinge to the largest value no further than 1.5 * IQR from the hinge (IQR = inter-quartile range, or distance between the first and third quartiles). The lower whisker extends from the hinge to the smallest value at most 1.5 * IQR of the hinge. Superimposed to each boxplot, the colored dots represent individual means. ** p < 0.01, *** p < 0.001. Uninf: uninformative, inf: informative, dis: distractor.

RT is shorter after low arousing music and uninformative cues when phasic arousal is increased.

Then, in order to test the effect of the arousing content of distractor in trials with a distractor, RT were fitted to a linear mixed model with Music, Cue, DisPosition2 (early or late dis) and DisContent (lowA or highA dis). As in the previous analysis, we found significant effects of the type of Cue (χ^2^ (1) = 47.16, p < 0.001, ω^2^⍰: very small) and the position of the distractor (χ^2^ (1) = 509.01, p < 0.001, ω^2^⍰: small), and a significant Music x Cue x DisPosition2 interaction (χ^2^ (1) = 4.08, p < 0.05, ω^2^⍰: very small).

Importantly, RT were significantly modulated by the arousing content of the distractor (χ^2^ (1) = 7.67, p < 0.01, ω^2^⍰: very small). RT were shorter after high arousing distractor (242.9 ± 1.5 ms) compared to low arousing distractor (252.3 ± 1.8 ms). A significant interaction DisContent x DisPosition2 was also observed (χ^2^ (1) = 6.18, p < 0.05, ω^2^⍰: very small). The DisContent effect was significant in the late (p < 0.01, d: very small), but not in the early (p = 0.91) dis condition, suggesting that the arousal content of the distracting sound influenced the RT only when the distractor was presented later during the delay (Fig. 4).

**Fig. 4.**
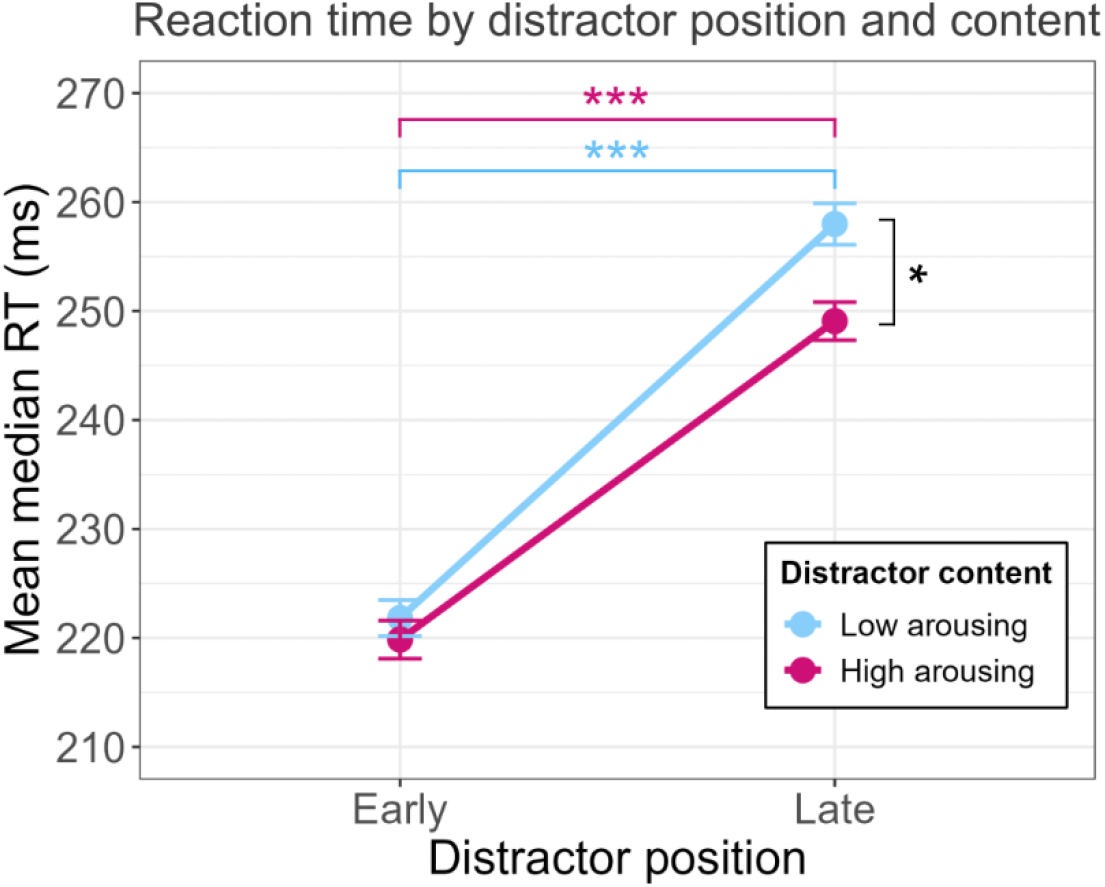
Mean RT by distractor position and arousing content. Each point corresponds to values averaged across subjects. Within-subject standard errors of the mean are plotted around group means. * p < 0.05, *** p < 0.001.

To investigate sustained attention, we explored the effect of the music type on RT variability (standard deviation of RT) in trials with no distractor. We found positive evidence against an effect of music (BF_10_ = 0.20) on RT variability.

#### 2.2.2. Music does not impact Response Type Rates

Participants performed well the task, with a high mean rate of hits (91.04 ± 1.25 %), and low mean rates of missed responses (0.66 ± 0.08 %), false alarms (8.28 ± 1.21 %), and late responses (0.03 ± 0.01 %). We found weak evidence for an effect of music on the hit rate (BF_10_ = 1.31), and moderate evidence against an effect of music on the miss rate (BF_10_ = 0.18), false alarms (BF_10_ = 0.24) and late responses (BF_10_ = 0.31). Response rates are not modulated by tonic arousal.

### 2.3. Music impacts Physiological measures of attention and phasic arousal

#### 2.3.1. PDR to the cue

To test the impact of music and cue types on phasic arousal to task-relevant stimuli, mean amplitudes of the PDR to the cue in trials with no distracting sound (Fig. 5) were analyzed in four time-windows following cue onset (500 – 1000 ms; 1000 – 1500 ms; 1500 – 2000 ms; 2000 – 2500 ms). No significant effect of music was found, and Bayesian paired-sample Wilcoxon signed-rank tests conducted for each time window confirmed weak to positive evidence for no effect of music (500 – 1000 ms: BF_10_ = 0.36; 1000 – 1500 ms: BF_10_ = 0.18; 1500 – 2000 ms: BF_10_ = 0.18; 2000 – 2500 ms: BF_10_ = 0.20). A significant effect of the cue on the PDR amplitude was found in the three time-windows from 500 to 2000 ms post-cue (χ^2^ (1) = 9.62, p < 0.01; ω^2^⍰]: medium; χ^2^ (1) = 21.23, p < 0.001; ω^2^⍰] = 0.17; χ^2^ (1) = 5.78, p < 0.05; ω^2^⍰]: small, respectively). In the absence of distracting sound, the PDR to the cue is larger after an informative cue than after an uninformative cue from 500 to 2000 ms, indicating increased phasic arousal with enhanced voluntary attention.

**Fig. 5.**
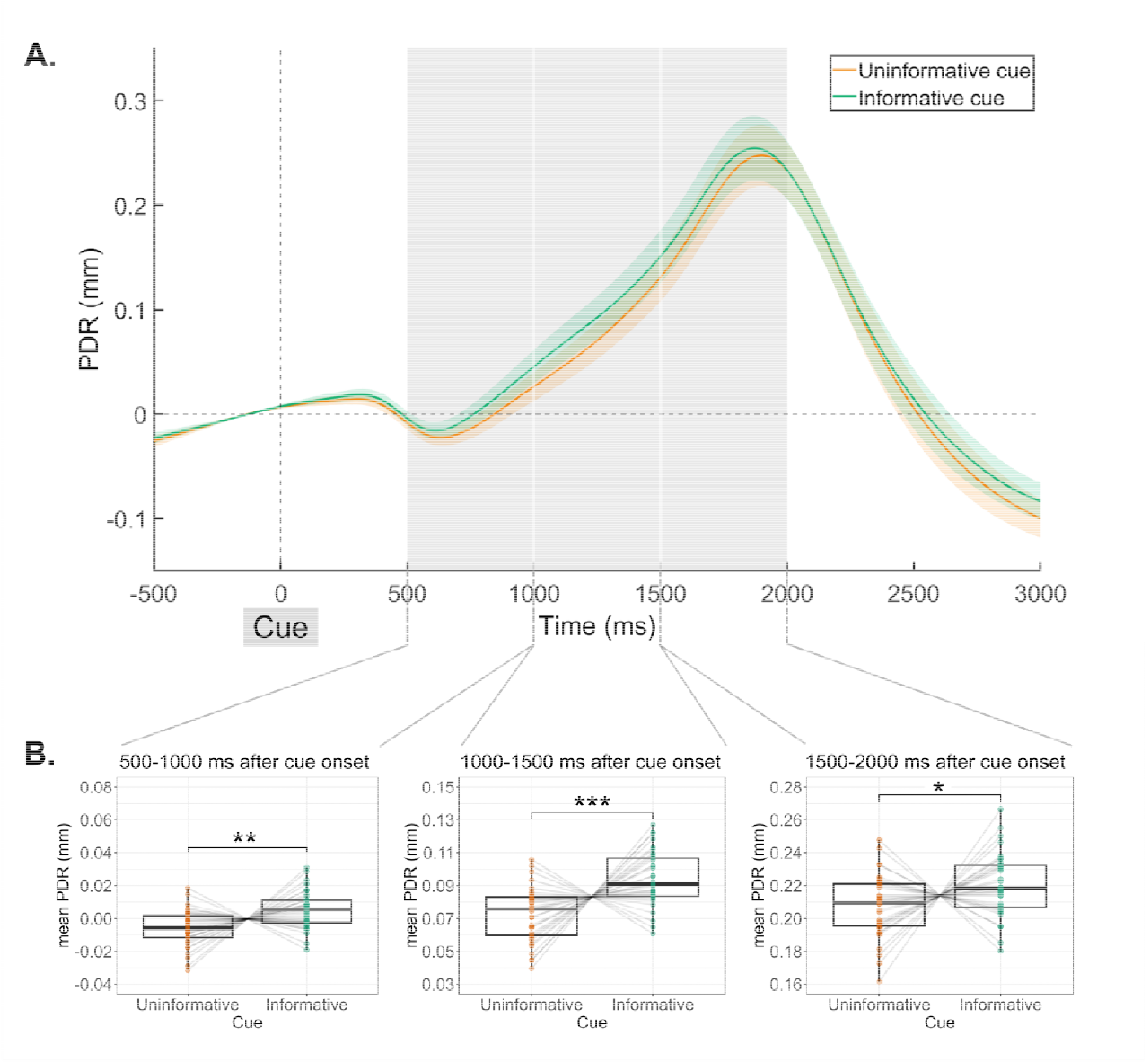
**Pupil Dilation Response (PDR) to the cue**, as a function of cue type. **A**. PDR time course. Shadowed areas represent standard errors of the mean. Grey areas correspond to time-windows where the cue effect (informative vs. uninformative) was significant. **B**. Mean values for the significant time-windows are presented below the PDR time-course. Boxplots were plotted using normalized individual data (= raw data – subject mean + group mean). Within each boxplot, the horizontal line represents the group median, the lower and upper hinges correspond to the first and third quartiles. The upper whisker extends from the hinge to the largest value no further than 1.5 * IQR from the hinge (IQR = inter-quartile range, or distance between the first and third quartiles). The lower whisker extends from the hinge to the smallest value at most 1.5 * IQR of the hinge. Superimposed to each boxplot, the colored dots represent individual means. * p < 0.05, ** p < 0.01, *** p < 0.001.

#### 2.3.2. PDR to the distractor

To test the impact of music on phasic arousal to salient task-irrelevant stimuli, mean amplitudes of the PDR to the distractor were analyzed in two time-windows following a distractor: 400 – 800 ms and 1000 – 1600 ms for early dis, and 400 – 800 ms and 1100 – 1700 ms for late dis.

During the first time-window, a significant main effect of the DisPosition2 (χ^2^ (1) = 4.73, p < 0.05; ω^2^⍰]: very small) and DisContent (χ^2^ (1) = 16.32, p < 0.001, ω^2^⍰]: small) was found on PDR amplitude. Specifically, the PDR was larger after an early (0.112 ± 0.005 mm) than after a late distractor (0.099 ± 0.005 mm), and larger after a high arousing (0.118 ± 0.005 mm) than after a low arousing distractor (0.093 ± 0.005 mm). Additionally, a significant interaction between DisPosition2 and DisContent (χ^2^ (1) = 4.49, p < 0.05; ω^2^⍰]: very small) was observed (Fig. 6). Post-hoc HSD analysis revealed that the PDR amplitude was larger in high arousing dis compared to low arousing dis in the late (p < 0.001; d: medium) but not in the early dis condition (p = 0.526).

**Fig. 6.**
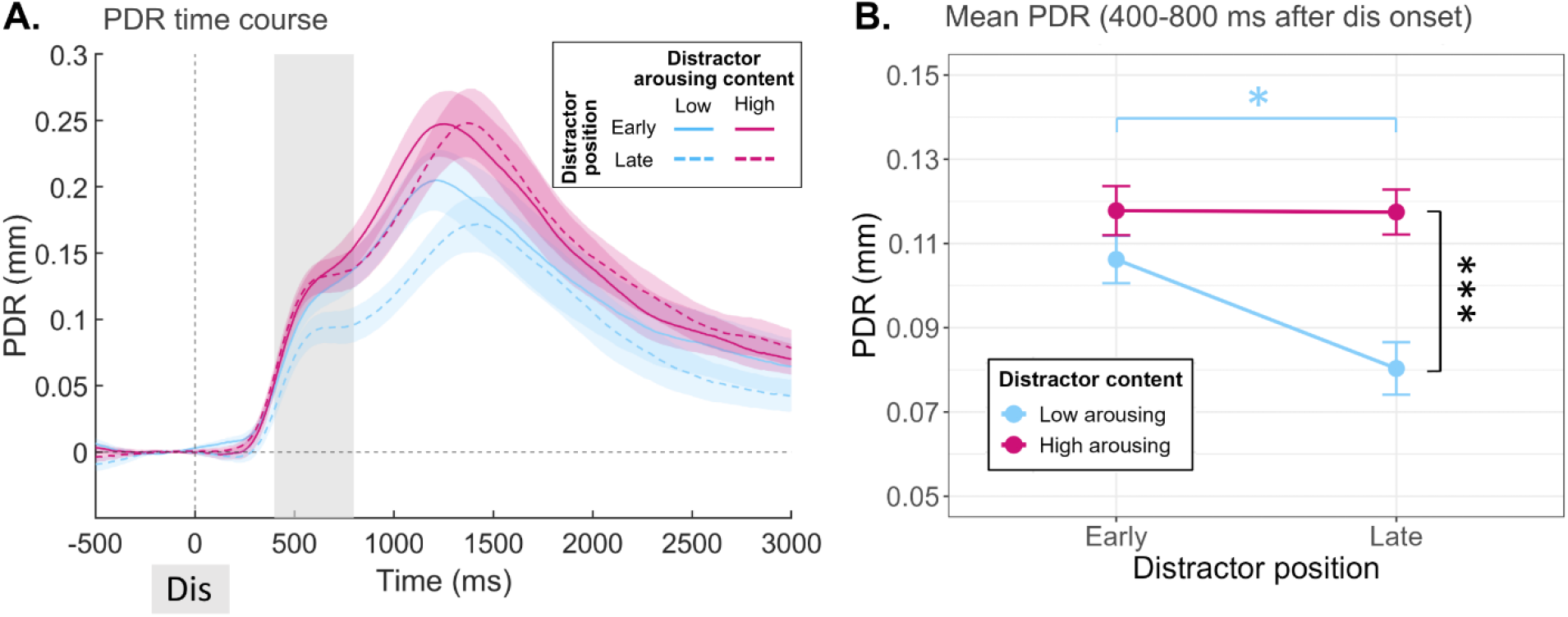
Pupil Dilation Response (PDR) to the presentation of the distractor sound, as a function of distractor position and content. **A**. PDR time-course. Shadowed areas represent standard errors of the mean. The grey area corresponds to the time window where the DisPosition x DisContent interaction was significant. **B**. Mean PDR amplitude in the 400 - 800 ms post-distractor. Each point corresponds to values averaged across subjects. Within-subject standard errors of the mean are plotted around group means. * p < 0.05, *** p < 0.001.

During the second time-window a significant main effect of the DisContent (χ^2^ (1) = 37.18, p < 0.001, ω^2^1]: medium) and a marginal effect of the DisPosition (χ^2^ (1) = 3.65, p = 0.056, ω^2^⍰]: medium) was found on PDR amplitude. Specifically, PDR was larger after a high arousing (0.228 ± 0.008 mm) than after a low arousing distractor (0.173 ± 0.008 mm) and tended to be larger after an early (0.209 ± 0.009 mm) than after a late distractor (0.192 ± 0.008 mm).

As already shown (Widmann et al., 2018), the PDR is larger after a high arousing rather than a low arousing distractor. Importantly, as observed for the RT, the effect of the arousal content on the PDR to the distractor occurred only when the distractor was presented later during the delay.

#### 2.3.3. ERP to the cue

To investigate the impact of music on brain correlates of voluntary attention orienting, we analyzed the Contingent Negative Variation (CNV) elicited between the cue and the target, in trials with no distracting sounds. Cue-related potentials were analyzed on *Cz* in two time-windows following the cue: 650 – 850 ms and 850 – 1050 ms. A significant effect of the Cue (χ^2^ (1) = 5.57, p < 0.05, ω^2^⍰: small) was found in the first, but not in the second time-windows and showed a larger CNV in the informative (−2.6 ± 0.2 µV) compared to uninformative (−2.2 ± 0.2 µV) cue condition (Sup. Fig. 1). The CNV is not affected by the type of music, as confirmed by Bayesian paired-sample Wilcoxon signed-rank tests conducted for each time window (650 - 850 ms: BF_10_ = 0.18; 850 - 1050 ms: BF_10_ = 0.19).

#### 2.3.4. ERP to the distractor

To investigate the impact of music on distractor processing, we investigated the successive typical ERPs to the distracting sounds: a fronto-central N1 (~110ms), a fronto-central early-P3 (~240ms) and a late-P3 with frontal and parietal components (~340ms).

The mean amplitude of N1 response to distractors was analyzed on **FCz** in the **90 – 130 ms** time-window following a distractor. No significant effect was found.

The mean amplitude of early-P3 response to distractors was analyzed on **Cz** in the **220 – 260 ms** time-window following a distractor. The early-P3 amplitude was found significantly modulated by DisContent (χ^2^ (1) = 55.96, p < 0.001, ω^2^1]: medium) and marginally by DisPosition2 (χ^2^ (1) = 3.57, p = 0.059, ω^2^1]: very small). Early-P3 amplitude was larger in response to a highly arousing distractor (5.58 ± 0.26 µV) compared to a low arousing distractor (3.40 ± 0.26 µV) and tended to be larger in response to a late distractor (4.77 ± 0.26 µV) compared to an early one (4.22 ± 0.27 µV). Importantly, significant interactions were also revealed: DisContent x DisPosition2 (χ^2^ (1) = 3.89, p < 0.05, ω^2^⍰]: very small) and DisContent x DisPosition2 x Music (χ^2^ (1) = 4.22, p < 0.05, ω^2^⍰]: very small). Post-Hoc HSD tests were conducted to analyze this triple interaction. Early-P3 amplitude was larger in response to a highly arousing distractor compared to a low arousing distractor in all conditions (p < 0.001, d: medium for low arousing music, early and late distractors, d: large for high arousing music, late distractors), except in trials with an early distractor after listening to high arousing music (p = 0.645) (Fig. 7). The impact of the distractor arousing content on the early-P3 to distractor is modulated by tonic arousal.

**Fig. 7.**
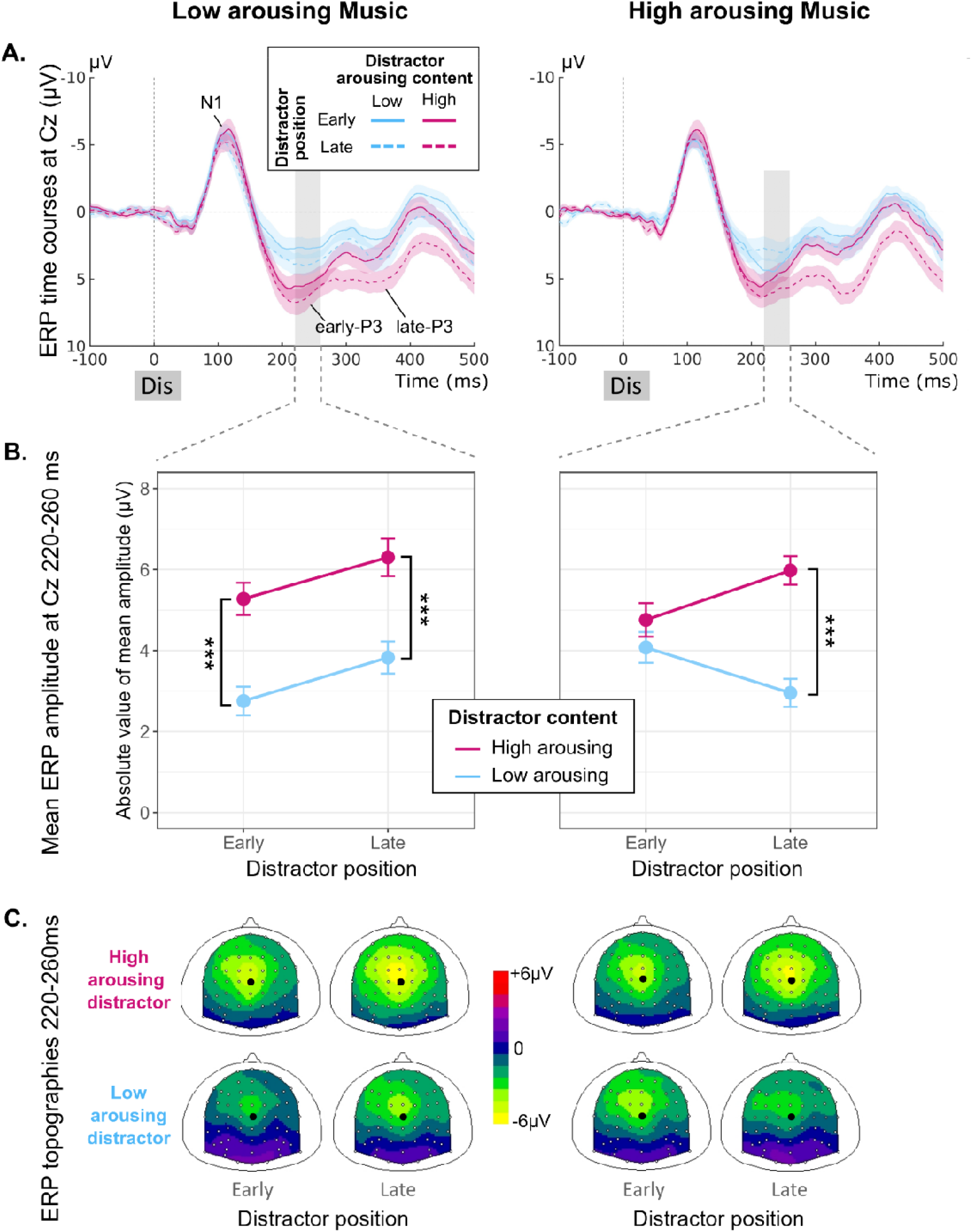
Event-Related Potentials (ERPs) on Cz electrode to the presentation of a distractor as a function of music type (left column: low arousing music; right column: high arousing music), distractor position (early vs late) and content (low arousing dis vs. high arousing dis). **A**. ERP time-course at Cz electrode. Shadowed areas represent standard errors of the mean. Grey areas correspond to the early-P3 time-window. **B**. ERP mean values for the early-P3 at Cz in the 220 – 260 ms time-window post distractor. **C**. Corresponding scalp topographies (top view). The larger black dot indicates the Cz electrode.

For the late-P3 response, the distractor-related potential was analyzed on **Fz** and **CPz** in the 320 – 360 ms time-window following a distractor. On **Fz**, a significant effect of DisPosition2 (χ^2^ (1) = 39.80, p < 0.001, ω^2^⍰]: medium) showed larger late-P3 amplitude after a late distractor (4.81 ±.26 µV) than after an early distractor (2.84 ± 0.27 µV). A significant effect of DisContent (χ^2^ (1) = 14.34, p < 0.001, ω^2^⍰]: small) showed larger late-P3 amplitude after a high arousing distractor (4.41 ± 0.26 µV) than after a low arousing distractor (3.23 ± 0.28 µV). A significant effect of Cue (χ^2^ (1) = 4.39, p < 0.05, ω^2^⍰]: very small) showed larger late-P3 amplitude in the informative (4.15 ± 0.24 µV) compared to uninformative cue condition (3.50 ± 0.29 µV). Additionally, a significant interaction between DisPosition2 and DisContent (χ^2^ (1) = 4.10, p < 0.05, ω^2^⍰]: very small) was observed. Post-hoc HSD analysis revealed that the amplitude was larger in high arousing dis compared to low arousing dis in the late (p <.001, d: medium) but not in the early dis condition (p = 0.599, Sup. Fig. 2).

On **CPz**, a significant effect of DisPosition2 (χ^2^ (1) = 20.34, p < 0.001, ω^2^⍰]: small) showed larger late-P3 amplitude after a late distractor (4.42 ± 0.28 µV) than after an early distractor (2.97 ± 0.28 µV). A significant effect of DisContent (χ^2^ (1) = 53.58, p < 0.001, ω^2^⍰]: medium) showed larger late-P3 amplitude after a high arousing distractor (4.87 ± 0.28 µV) than after a low arousing distractor (2.52 ± 0.28 µV).

#### 2.3.5. ERP to the target

To test the impact of music and cue types on target processing, we analyzed the successive fronto-central N1 (~130ms) and parietal target-P3 (200-500ms), in trials with no distracting sound.

The mean amplitude of N1 response to targets was analyzed on **FCz** in the **110 – 150 ms** time-window following a target. A significant effect of Cue (χ^2^ (1) = 5.88, p < 0.05, ω^2^⍰]: small) indicated a smaller N1 amplitude after an informative (−5.46 ± 0.22 µV) compared to an uninformative cue (−5.81 ± 0.25 µV; Sup. Fig. 3).

The mean amplitude of the target-P3 response was also analyzed on **CPz** in the **250 – 500 ms** time-window following a target. A significant effect of cue (χ^2^ (1) = 23.56, p < 0.001, ω^2^⍰]: large) indicated a smaller target-P3 amplitude after an informative (7.87 ± 0.31 µV) compared to an uninformative cue (8.65 ± 0.34 µV; Sup. Fig. 3).

These data show reduced target processing after informative cues.

## 3. DISCUSSION

The present study aimed to explore how tonic and phasic arousal influence voluntary and involuntary attention and shape performance, in the auditory modality. To do so, we used the Competitive Attention Task (CAT), a paradigm that distinguishes voluntary attention through cue informativeness and involuntary attention through the presence of unexpected task-irrelevant salient sounds, called distractors.

### 3.1. Increasing phasic arousal shortens reaction times

In this study, several parameters shorten reaction times. First, informative cues, compared to uninformative ones (cue effect), lead to shorter reaction times, consistent with previous findings (Fig. 8.A, Bidet-Caulet et al., 2015; ElShafei et al., 2018). A cue effect is also observed on physiological measures, with a larger pupil dilation and an enhanced Contingent Negative Variation (CNV) prior to target onset, as well as a reduced P3 in response to the target, in the informative compared to the uninformative cue condition. These physiological effects, nicely mirroring the behavioral results, are consistent with previous results (Bidet-Caulet et al., 2015; Grandjean et al., 2024). Pupil dilation is commonly associated with arousal (Gilzenrat et al., 2010; Joshi et al., 2016; Nieuwenhuis et al., 2011), the CNV reflects anticipatory attention (Brunia & van Boxtel, 2001) and the P3 is linked to late-stage processing of relevant stimuli (Duncan-Johnson & Donchin, 1982; Mars et al., 2008; Suwazono et al., 2000). Cue informativeness thus triggers engagement of voluntary attention, which modulates phasic arousal responses (Fig. 8.C). We therefore demonstrate that the combined enhancement of voluntary anticipatory attention and phasic arousal mechanisms following informative cues leads to optimized target processing and faster target detection.

**Fig. 8.**
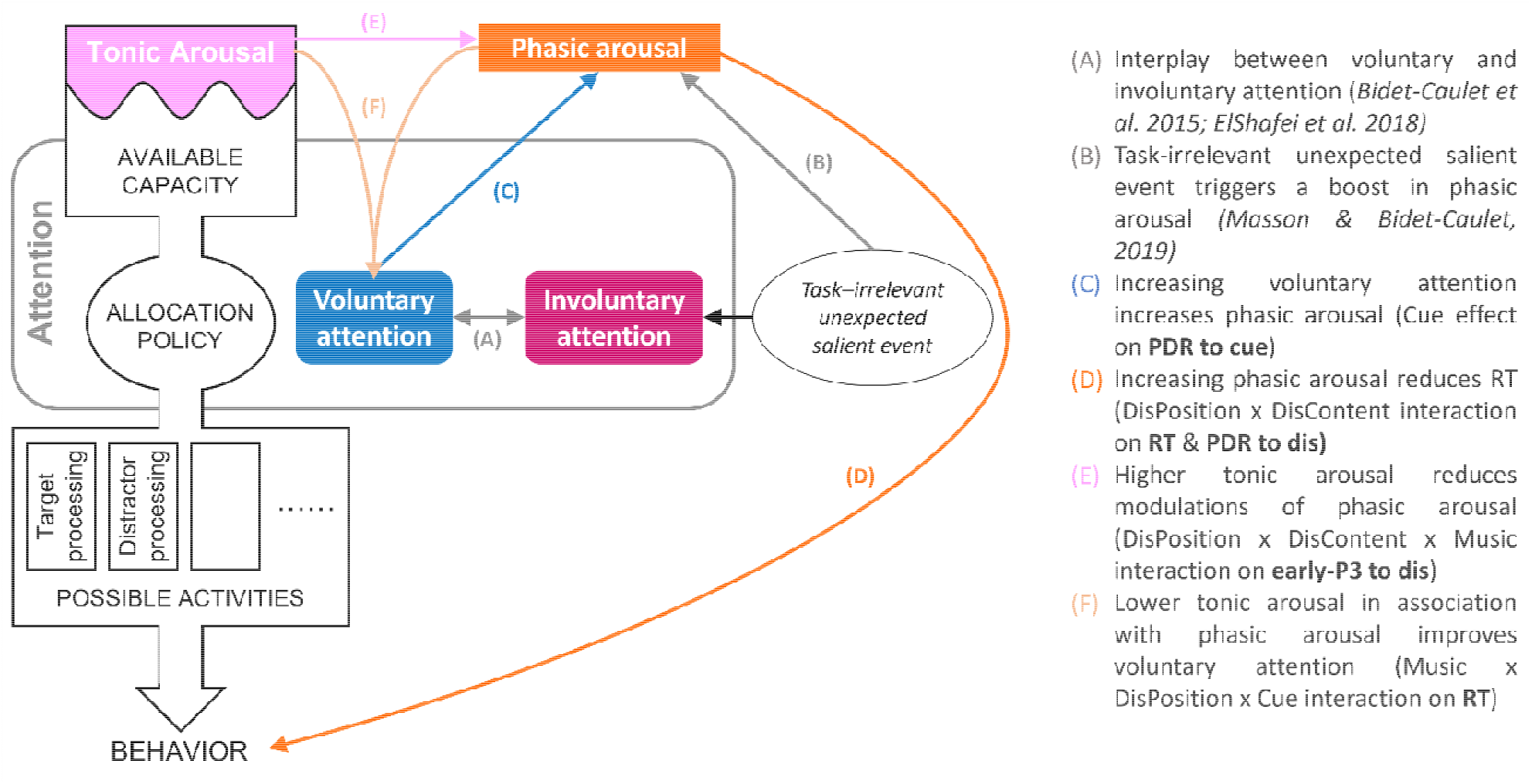
Schematic illustration of arousal and attention interactions (after Kahneman, 1973). This model reflects a general framework for viewing how arousal and attention processes could interact. Overall, the level of tonic arousal determines the amount of available capacity for attention, and attention allocates these resources to different possible activities resulting in an adapted behavior. As shown by previous studies using the CAT, a balance between involuntary and voluntary attention (A) controls the allocation of available capacity to the possible brain activities, task-relevant ones (e.g. target processing) or task-irrelevant ones (e.g. distractor processing). Task-irrelevant unexpected salient events (distractors) trigger both an increase in phasic arousal (B) and involuntary attention processes. In the present study, we showed that voluntary attention increases phasic arousal (C), which can in turn impact behavior (D). The level of tonic arousal can also influence phasic arousal (E). The interaction between tonic and phasic arousal allows to reach an optimal performance, possibly by improving anticipatory mechanisms (F).

Second, challenging the conventional definition of distraction, early distractors —occurring well before target onset— can actually reduce reaction times, in comparison to trials with late or no distractors, as previously observed with the CAT (Bidet-Caulet et al., 2015; Hoyer et al., 2021). This facilitation effect has been attributed to a boost in phasic arousal (Fig. 8.B, Masson & Bidet-Caulet, 2019). It has been proposed that distractors elicit two distinct phenomena: a brief involuntary orienting of attention (so-called distraction) and a more sustained increase in phasic arousal. The final behavioral outcome depends on the balance between the costs of distraction and the benefits of phasic arousal. A large pupil dilation response is observed in response to distractors, confirming an increase in phasic arousal after unexpected task-irrelevant salient sounds. At the cerebral level, event-related potentials (ERPs) to distractors are used as markers of distractor processing (Escera et al., 2000). However, these ERPs encompass both the involuntary orienting of attention and the phasic arousal responses triggered by distractors (Masson & Bidet-Caulet, 2019). The early-P3 – peaking around 240ms after distractor onset at fronto-central electrodes – has been proposed to reflect the arousing properties of distracting sounds (Masson & Bidet-Caulet, 2019).

Finally, increasing the arousing content of distractors results in shorter reaction times. In agreement with previous works (Masson & Bidet-Caulet, 2019; Widmann et al., 2018), high arousing distractors elicit larger pupil dilation responses and greater early-P3 amplitudes than low arousing distractors, indicating a larger phasic arousal response. To our knowledge, this is the first evidence that such physiological effects translate into measurable behavioral outcomes, demonstrating a clear performance benefit associated with the arousing content of distractors (Fig. 8.D).

In summary, both cue- and distractor-related facilitation effects on reaction times reported in the literature are replicated in the present data. Importantly, the multimodal approach used in the present study further demonstrates that these behavioral benefits are associated to phasic arousal —as reflected by pupil dilation responses and early-P3 amplitudes— or voluntary attention processes, indexed by the CNV. Importantly, the observed effect of distractor arousing content on reaction times supports the hypothesis that the facilitation effect of unexpected task-irrelevant salient sounds is driven by an increase in phasic arousal, as proposed by Masson and Bidet-Caulet, 2019.

### 3.2. Tonic and phasic arousal jointly modulate behavior

First of all, we confirmed that the arousal manipulation with music was effective: high – compared to low – arousing music excerpts resulted in larger pupil size and faster heart rate, indicating elevated tonic activity in the locus coeruleus-norepinephrine (LC-NE) system. We next investigated how tonic arousal influences the behavioral and physiological dynamics of attentional control.

Importantly, we demonstrate that the tonic arousal level impact reaction times. In the low arousing music condition, the cue effect is absent following an early distractor. Notably, this absence is driven by a reduction in reaction times in the uninformative cue condition rather than by an increase in reaction times in the informative cue condition, when comparing low to high arousing music contexts. Thus, lower tonic arousal level appears to be beneficial only when the cue fails to provide enough information to effectively guide and adjust their anticipation, and when associated with an increase in phasic arousal. Specifically, low arousing music induces a tonic arousal state that may improve behavior by fostering a cognitive state more conducive to anticipatory mechanisms (Fig. 8.F). This facilitatory effect seems to be absent after a late distractor, presumably due to the limited time available before target onset for the effect to manifest. These data demonstrate that, in a simple detection task, the combination of tonic and phasic arousal effects allows to reach an optimal level of arousal, which in turn leads to optimal performance, while requiring less engagement of voluntary attention resources.

This finding aligns with studies reporting improved performance under lower tonic arousal states, whether induced by isometric exercise (Mather et al., 2020) or music (Kuan et al., 2018; Nadon et al., 2021). The effect of music-induced modulations is also consistent with the arousal–mood hypothesis, which proposes that slow tempo/minor-mode (low arousing) music can improve cognitive control and processing speed under certain conditions (Quan et al., 2023). Interestingly, the low and high arousing music excerpts selected in the present study did not differ in tempo, suggesting that the observed effect are specifically linked to arousal. Conversely, other studies have found that increasing arousal through music can improve performance (Cloutier et al., 2020; Kiss & Linnell, 2021), likely by helping participants reach an intermediate, optimal level of arousal on the inverted U-shaped curve relating arousal to performance (Aston-Jones & Cohen, 2005). These discrepancies may be explained by differences in task demands, with more monotonous tasks benefiting from increase in tonic arousal, and by individual differences in baseline arousal levels.

### 3.3. Phasic arousal and voluntary attention jointly modulate behavior

Interestingly, the effect of distractor arousing content is modulated by the distractor’s position in the cue-target interval, both behaviorally and physiologically. The beneficial effect of high arousing content on reaction times emerges only when the distractors occur late in the interval. In contrast, early distractors speed up responses regardless of their arousing content. A mirror pattern is observed on the pupil dilation response: early distractors elicit a larger PDR irrespective of their arousing content, whereas a reduced PDR is seen specifically for late low arousing distractors.

One interpretation is that the probability of a sound appearing gradually increases during the delay between the cue and the target (Coull & Nobre, 1998; Griffin et al., 2001; Niemi & Näätänen, 1981). As a result, early distractors are more surprising and evoke a strong phasic arousal response, reflected in a large PDR and shorter reaction times, regardless of the distractor’s arousing content. In contrast, late distractors are less surprising and elicit a weaker arousal response which can be modulated by the arousing content. In addition to this effect of surprise, the gradual build-up of voluntary attention during the cue–target interval (Coull & Nobre, 1998; Griffin et al., 2001; Niemi & Näätänen, 1981; Posner et al., 1980) may also shape the processing of distractors. This gradual build-up is confirmed by the progressive increase in CNV amplitude during the cue–target interval, with a significant cue effect between 650 and 850 ms post cue onset. Thus, early distractors occur before voluntary attention is fully established, and their simple occurrence is sufficient to trigger a strong phasic response. In contrast, when more voluntary selective attention is engaged later in the interval, the processing of task-irrelevant stimuli may be dampened, resulting in a reduced PDR in particular to low arousing distractors. Finally, a floor effect on reaction times in the early distractor condition may limit the potential for further speed gains due to arousing content. By contrast, the generally slower responses following late distractors leave more room for high arousing stimuli to counteract distraction costs.

These data suggest that the effects of distractors on behavior depend on the dynamic interplay between phasic arousal and voluntary attention over time.

### 3.4. Tonic and phasic arousal shape the cortical responses to distractors

When considering the early-P3 cortical response elicited by the distractor, the interaction between distractor position and arousing content is further modulated by the level of tonic arousal resulting from the musical induction. On the one hand, following high arousing music, we observe a pattern that closely mirrors reaction time and PDR results: in late position only, a highly arousing distractor produces a larger early-P3 than a low arousing one. On the other hand, under lower tonic arousal, high arousing distractors evoke larger early-P3 amplitudes than low arousing ones in both early and late positions.

One possible explanation is that high arousing music may raise tonic arousal to a level where phasic arousal responses are saturated, especially for early distractors that are more surprising. In this case, the modulation by the distractor arousing content may be overshadowed, resulting in similarly large early-P3 amplitudes for both low and high arousing distractors in the early position. Alternatively, after low arousing music, the level of tonic arousal is lower and phasic arousal responses can be modulated by the distractor arousing content in both early and late positions. This pattern is in line with models of the LC-NE system, which propose that high tonic arousal can saturate phasic responses, limiting further modulation by stimulus salience especially when phasic responses are already strong (Aston-Jones & Cohen, 2005). This is in line with the work of Mather et al., 2020, who used different method and protocol, but similarly, found that reducing tonic arousal increased phasic arousal responses. This effect of tonic arousal is not observed on RT nor PDR, possibly because these measures are less sensitive to subtle or short-lasting effects than ERPs. Indeed, the early-P3 is measured within a narrow 220–260⍰ms window after distractor onset, whereas PDR effects emerge later, between 400–800⍰ms, and reaction times reflect the combined influence of all preceding processes.

Therefore, the present data show that increasing tonic arousal can saturate phasic arousal responses at the cortical level (Fig. 8.E), in line with an inverted U-shaped relationship between tonic and phasic arousal in humans.

#### General conclusion

Using a multimodal approach, this study demonstrates that attention and arousal jointly shape behavioral performance. First, we provide empirical findings confirming the hypothesis that the behavioral beneficial effect of distractors, reported in the literature, is associated with a burst of phasic arousal, with faster reaction times associated with larger pupil dilation responses. Second, we reveal an interplay between phasic arousal and voluntary attention, where an increase in phasic arousal can substitute for voluntary attention engagement, possibly by relieving attentional resources. Finally, we show that tonic arousal, modulated by music, interacts with phasic arousal to approach an optimal arousal level. This interaction can boost performance by preserving phasic responsiveness under low tonic arousal or lead to phasic saturation at physiological level when tonic arousal is too high. Together, these results are in line with an inverted U-shaped relationship between arousal and performance, highlighting the importance of adjusting tonic and phasic arousal for optimal attentional control.

## 4. MATERIAL AND METHODS

### 4.1. Participants

Forty participants took part in the experiment. The data of six participants had to be discarded because of technical issues or noisy data. Then, the analysis was performed on data from thirty-four participants (17 female, 5 left-handed and 29 right-handed, mean age ± standard deviation (SD): 23.15 ± 2.66 years). All participants were free from neurological or psychiatric disorder, did not take any medication that could impact brain functioning, and had normal hearing and normal or corrected-to-normal vision. The study was approved by a national ethical committee, and subjects gave written informed consent, according to the Declaration of Helsinki, and they were paid for their participation. To limit inter-individual differences in tonic arousal, only participants drinking no more than two cups of coffee (or equivalent in caffeinated drinks) and one glass of alcohol per day were included.

The sample size was determined based on common practices for similar neurophysiological studies (Bidet-Caulet et al., 2015; Dragone et al., 2018; ElShafei et al., 2018; Steiner & Barry, 2011; Wang et al., 2018; Wetzel et al., 2016) and was maximized as much as possible given the available resources.

### 4.2. Stimuli and Task

The task consisted of twenty-four blocks (separated in two sessions of twelve blocks each, one to six days apart, in the morning). Each block started with a music track played during 30s followed by a 2s pause and then by 48 trials of the Competitive Attention Task (CAT; Fig. 1). To modulate tonic arousal, this musical excerpt was either low or high arousing (*lowA* or *highA* music) (see Supplementary Materials for more details).

Each trial of the CAT started with a visual cue (200ms duration), corresponding to a green arrow pointing either to the left, to the right or to both sides, centrally presented on a grey background screen. The visual cue was followed by a delay randomly chosen between 900 and 1100ms. Then, a target sound (50ms duration), a monaural harmonic sound (fundamental frequency: 200 Hz, 5 harmonics; 5ms rise-time, 5ms fall-time), was presented in earphones. In 25% of the CAT trials, a binaural distracting sound (300ms duration) was played during the delay between the cue and the target. In the absence of the visual cue, a blue fixation cross was presented at the center of the screen.

The cue and target categories were manipulated in the same proportion for trials with and without distracting sound. In 66.6% of trials, the cue was informative (pointing left and the target sound was played in the left ear (33.3%), or pointing right and the target sound was played in the right ear (33.3%)). In the last 33.3% of the trials, the cue was uninformative, pointing in both directions while the target sound was played in the left (16.7%) or right (16.7%) ear.

A total of 24 different sounds were used as distracting sounds (clock-alarm, doorbell, laughter, gunfire, etc.). They were chosen from a pool of sounds from Bidet-Caulet et al. (2015) and Widmann et al. (2018) in order to modulate phasic arousal: 12 low arousing and emotionally positive distracting sounds (*lowA dis*), and 12 high arousing and emotionally negative distracting sounds (*highA dis*) (see Supplementary Materials for more details). These distracting sounds were binaurally presented, and their onset equiprobably fell in two different time periods before the target onset: in the 600-800ms range (*early dis*), or in the 350-550ms range (*late dis*). To compare behavioral and physiological responses to acoustically matched sounds, the same distracting sounds were played for each combination of music type (lowA and highA music), cue category (informative left, informative right and uninformative) and distractor condition (early or late dis). Each distracting sound was played 12 times during the whole experiment, but no more than twice during each single block to limit habituation.

Participants were instructed to perform a detection task by pressing a mouse button as fast as possible when they heard the target sound, and to ignore the distracting sounds. They were asked to focus their attention to the cued side in the case of informative cues. Moreover, they were informed that informative cues were 100% predictive and that a distracting sound could be sometimes played.

Participants were also instructed to blink naturally during the task but to keep their eyes fixated on the cross and to minimize eye movements while performing the task. Participants had between 3000 and 3200ms to answer after target sounds, each CAT trial lasted therefore around 4350ms on average, leading to block duration of ≈4 min and EEG session of ≈1h (breaks included).

Participants performed a total of twenty-four blocks (48 trials each). Within blocks, CAT trials were randomized differently for each subject to limit sequence effects. In each subject, across both sessions, the same twelve blocks were presented in the low and in the high arousing music conditions. In each session, high and low arousing blocks were alternating, with one session starting with a low arousing music and the other one with a high arousing music. The order of sessions was balanced across participants.

During the whole experiment, for each music condition, 144 no dis, 12 early lowA dis, 12 early highA dis, 12 late lowA dis, 12 late highA dis were presented in each cue category (informative left, informative right & uninformative).

### 4.3. Procedure

For each session, participants were seated in a comfortable armchair at a 1 m distance from the screen. All stimuli were delivered using Presentation® software (Version 23.0, Neurobehavioral Systems, Inc., Berkeley, CA, www.neurobs.com). Sounds were delivered through earphones (Etymotic, ER4SR). First, the auditory threshold was determined for the target sound, in each ear, for each participant using the Bekesy tracking method (Haupt, 2003). The target sounds were monaurally presented at 10 dB SL (between 23 and 31 dBA across subjects), while distracting sounds were binaurally played at 40 dB SL (between 69 and 77 dBA across subjects), low arousing music at 40 dB SL (between 54 and 62 dBA across subjects), high arousing music at 45 dB SL (between 69 and 77 dBA across subjects) above the target sound thresholds. Second, participants were trained with a short sequence of the task. Finally, for each session, the electroencephalogram (EEG), the electrooculogram (EOG), the electrocardiogram (ECG), the skin conductance and the pupil size were recorded while participants performed 12 blocks. Each block started with a five-point eye-tracker calibration and validation procedure.

### 4.4. Data recordings

#### 4.4.1. Physiological recordings

EEG data were recorded from 64 electrodes using the ActiveTwo system (BioSemi, the Netherlands). Eight additional Flat-Type active electrodes were used for vertical EOG (left supraorbital and infraorbital ridge locations), ECG (left shoulder and stomach), and skin conductance (medial phalanx of index and middle fingers of the non-dominant hand) recordings, and two electrodes were placed on earlobes for offline referencing. Data were amplified (−3 dB at ~204 Hz low-pass, DC coupled), digitized (1024 Hz), and stored for offline analysis. EEG data were re-referenced offline to the average potential of the two earlobe electrodes.

#### 4.4.2. Pupillometry

The pupil size of the left eye was recorded during each entire block and monitored using an eye-tracking system (Eyelink Portable Duo, SR Research Ltd., Mississauga, Ontario, Canada). Participants were placed at a 45 cm distance from the eye-tracker (remote mode, sampling rate of 500 Hz). At the beginning of each block, a five-point calibration followed by a five-point validation procedure was performed.

### 4.5. Data analysis

Data analysis was performed using the software package for electrophysiological analysis (ELAN Pack, Aguera et al., 2011) developed at the Lyon Neuroscience Research Center and custom MATLAB programs.

#### 4.5.1. Behavior

For each participant, the shortest reaction time (RT) limit for correct response (RT lower limit) was set to 150 ms after the target, from previous studies using a similar paradigm in adults (Hoyer, Abdoun, et al., 2023; Hoyer et al., 2021; Hoyer, Pakulak, et al., 2023). The longest RT for a correct response (RT upper limit) was calculated from all RT > 0 ms using the John Tukey’s method of leveraging the Interquartile Range (3^rd^ Quartile + 1.5 x Interquartile Range). A button press before the RT lower limit was considered as a false alarm. A trial with a button press after the RT upper limit was considered as a late response. A trial with no button press before the next cue onset was considered as a missed trial. A trial with no FA and with a button press between the RT lower limit and upper limit was counted as a correct trial. For statistical analysis of reaction times (RT), all RT > 0 ms were considered. Additionally, the standard deviation of positive reaction time in trials with no distractor was calculated for each subject.

#### 4.5.2. Skin conductance

Skin conductance data were low-pass filtered offline at 10Hz. For each block, a Z-score normalization to the baseline (period of 2500ms before the onset of the music) was applied.

#### 4.5.3. Heart rate

ECG data were filtered offline with a 1Hz high-pass filter and a 40Hz low-pass filter. During the 30s music presentation, R-peaks were automatically detected, interbeat interval (IBI) were derived from the signal, and artifacts and ectopic beats were hand corrected using the QRSTool software (Allen et al., 2007). Then, the heart rate was extracted using the CMetX software (Allen et al., 2007) (www.psychofizz.org). Analyses were conducted on mean heart rate values.

#### 4.5.4. Pupillometry

The eye tracker pupil size digital counts were converted to millimeters using an artificial eye. Full blinks were detected and marked by the eye tracker. Partial blinks were detected from the smoothed velocity time as pupil size changes exceeding a velocity adjusted for each subject around 15 mm/s. Data were removed in a 50ms pre-blink and a 100ms post-blink interval and a 50ms pre-partial blink and 150ms post-partial blink interval. Subsequently, segments with blinks or missing data were interpolated with linear interpolation. Trials with more than 50% of interpolated data were removed from further analysis. Three distinct analyses were conducted with pupillometry data: the pupil size changes during the music presentation, the pupil dilation responses (PDR) to the cue, and the PDR to the distractor.

To investigate the effect of music presentation, a baseline subtraction was applied (period of 455ms before the onset of the music) and the mean amplitude was computed during the 30s music presentation.

To analyze the PDR to the cue, a baseline subtraction was applied using the 250ms period before stimulus onset and the mean PDR was computed in 500ms time-windows from 500 to 2500ms after cue onset. To analyze the PDR to the distractor, for each distractor onset time-range, surrogate dis-locked PDR were created in the no dis trials and subtracted from the actual dis-locked responses. The obtained PDR was thus clear of cue-related activity. Additionally, a baseline subtraction was applied using the 250ms period before stimulus onset and the mean PDR was computed in time-windows after distractor onset: 400 – 800 ms and 1000 – 1600ms post early dis, and 400 – 800 ms and 1100 – 1700 ms post late dis.

#### 4.4.5. EEG

EEG data were filtered offline with a 0.1 Hz high-pass filter and a 40 Hz lowpass filter. Eye-related activities were detected using independent component analysis (ICA) and were selectively removed via the inverse ICA transformation (Delorme & Makeig, 2004) (EEGLAB toolbox). Only 1 or 2 ICs were removed in each participant. In 21 participants, the flat or excessively noisy signals at 1 to 6 electrodes were replaced by their values interpolated from the remaining electrodes using spherical spline interpolation(Perrin et al., 1989). Trials including false alarms or undetected target, and trials contaminated with excessive muscular activity were excluded from further analysis.

ERPs were averaged for each stimulus event: cue-related potentials (cueRPs) were averaged locked to cue onset, target-related potentials (targetRPs) were averaged locked to target onset, and distractor-related potentials (disRPs) were averaged locked to distractor onset. Different baseline corrections were applied according to the investigated processes. ERP scalp topographies were computed using spherical spline interpolation (Perrin et al., 1989).

To investigate the deployment of top-down attention mechanisms in the absence of distracting sound (no dis), cueRPs were baseline corrected to the mean amplitude of the −100 to 0ms period before cue onset, and targetRPs were corrected to the mean amplitude of the −100 to 0ms period before target onset. To analyze ERPs to distracting sound, for each distractor onset time-range, surrogate disRPs were created in the no dis trials and subtracted from the actual disRPs. Then, disRPs were baseline corrected to the mean amplitude of the −100 to 0ms period before distractor onset. The obtained disRPs were thus clear of cue-related activity.

For the analysis of EEG and pupil data, we considered only trials that were not rejected in both EEG and pupil data analysis. In the low arousing music condition, the average number of trials per participant included in the analysis was: 267 informative cue no dis trials (Standard error of the mean, SEM = 3.3), 136 uninformative cue no dis trials (SEM = 1.3), 32 early lowA dis trials (SEM = 0.6), 31 early highA dis trials (SEM = 0.8), 33 late lowA dis trials (SEM = 0.4), and 33 late highA dis trials (SEM = 0.5). In the high arousing music condition, the average number of considered trials per participant was similar: 268 informative cue no dis trials (SEM = 3), 136 uninformative cue no dis trials (SEM = 1.4), 32 early lowA dis trials (SEM = 0.7), 31 early highA dis trials (SEM = 0.8), 33 late lowA dis trials (SEM = 0.5), and 33 late highA dis trials (SEM = 0.5).

### 4.6. Statistical analysis

When only two paired conditions were compared, Bayesian non-parametric Wilcoxon signed-rank test were conducted using JASP software (JASP—A Fresh Way to Do Statistics, 2021; Version 0.14.1). In contrast to Frequentist statistics, Bayesian analyses allow one to assess the credibility of both the alternative and null hypotheses. We reported Bayes factor (BF_10_) as a measure of evidence in favor of the null hypothesis (BF_10_ 0.33–1, 0.1–0.33, 0.01–0.1, and lower than 0.01: weak, moderate, strong, and decisive evidence, respectively) and in favor of the alternative hypothesis (value of 1–3, 3–10, 10–100, and more than 100: weak, moderate, strong, and decisive evidence, respectively (Lee & Wagenmakers, 2014)

When the analysis included more than one factor, we conducted linear mixed models (LMM) using R (R Core Team, 2021, version 4.1.2). We estimated the effect of the manipulated task parameters (Music, Cue, Distractor Position, Distractor Content) on measured variables (reactions times, pupil dilation responses, event-related potentials; see Table 1 for a list of the model characteristics). We accounted for the heterogeneity between-subjects and experimental conditions by defining them as effects with a random intercept, thus instructing the model to correct for any systematic differences in variability between participants and conditions. Values of the marginal R^2^ and conditional R^2^ (when available) can be found in Table 1. We ran a type II analysis of variance on the selected models. Wald chi-square tests were used for fixed effects in linear mixed-effects models. Frequentist models and statistics were performed using the lme4 (Bates et al., 2015) and car (Fox & Weisberg, 2019) packages. We considered the results of all analyses significant at p < 0.05. When we found a significant main effect or interaction, posthoc Tukey HSD tests were systematically performed using the emmeans package. P-values were considered as significant at p < 0.05 and were adjusted for the number of performed comparisons.

To compute effect sizes for the LMM effects, omega squared values were obtained using the R effectsize package and interpreted using Field’s (2013) guidelines: ω^2^⍰ < 0.01 = very small, 0.01 ≤ ω^2^⍰ < 0.06 = small, 0.06 ≤ ω^2^⍰ < 0.14 = medium, and ω^2^⍰ ≥ 0.14 = large. For post-hoc comparisons, Cohen’s d values were calculated with the R emmeans package and interpreted using Cohen’s (1992) criteria: *d* < 0.2 = very small, 0.2 ≤ *d* < 0.5 = small, 0.5 ≤ *d* < 0.8 = medium, and *d* ≥ 0.8 = large.

When no effect of music was found using LMMs, we conducted Bayesian Wilcoxon signed-rank tests to assess whether the null hypothesis was supported.

#### 4.6.1. Peripheral indices of arousal

Non-parametric statistics were used to test the effect of music on skin conductance, pupil size and heart rate. A paired Wilcoxon signed-rank test was used on the skin conductance mean amplitude, mean pupil size and mean heart rate during the presentation of the music (0-30 sec) to test the tonic arousal modulation.

#### 4.6.2. Behavior

##### Response Type Rates

The effect of music on the rate of Hits, Miss, False Alarms, and Late Responses was investigated using Bayesian non-parametric paired Wilcoxon signed-rank tests.

##### Reaction Times

Raw RT were log-transformed at the single-trial scale to be better fitted to a linear model with Gaussian family distribution. In a first model, logRT were fitted to a model with the following three within-subject fixed-factors: Music (2 levels: highA music, lowA music), Cue (2 levels: informative, uninformative) and DisPosition3 (3 levels: no dis, early, late). In a second model, to analyse the effect of the distractor content, logRT in trials with a distractor were fitted to a model with the following four within-subject fixed-factors: Music (2 levels: highA music, lowA music), Cue (2 levels: informative, uninformative), DisPosition (2 levels: early, late), DisContent (2 levels: highA dis, lowA dis).

Additionally, we investigated the effect of music type on the variability of reaction times using a Bayesian paired-sample Wilcoxon signed-rank test.

#### 4.6.3. Pupil dilation response (PDR)

For statistical analysis of the PDR to the cue (no dis trials), the mean pupil size per subject in the 500- 1000ms, 1000-1500ms, 1500-2000ms and 2000-2500ms time windows following a cue was fitted to a linear model with Gaussian family distribution model with two within-subject fixed-factors: Music (2 levels: highA music, lowA music) and Cue (2 levels: informative, uninformative).

For statistical analysis of the PDR to the distractor, mean pupil size was calculated separately for early and late distractors: in the 300-800ms and 950-1450ms time-windows following an early distractor, and in the 300-800ms and 1100-1600ms time-windows following a late distractor. Then, mean amplitude for the first and second time-windows were fitted to a linear model with Gaussian family distribution model. with four within-subject fixed-factors: Music (2 levels: highA music, lowA music), Cue (2 levels: informative, uninformative), DisPosition2 (2 levels: early, late), DisContent (2 levels: highA dis, lowA dis).

#### 4.6.4. Event-related potential (ERP)

Statistical analyses were performed on specific electrode and time-windows for each main ERP component (N1, early-P3 and late-P3). The electrodes and time-windows were chosen based on previous studies (Bidet-Caulet et al., 2015; Masson & Bidet-Caulet, 2019).

For statistical analysis of cue-related potentials (no dis trials), the mean amplitude per subject of the Contingent Negative Variation (CNV) on Cz electrode in the 650-850 ms and 850-1050 ms time-windows following a cue was fitted to a linear model with Gaussian family distribution model with two within-subject fixed-factors: Music (2 levels: highA music, lowA music) and Cue (2 levels: informative, uninformative).

For statistical analysis of distractor-related potentials, the mean amplitude per subject of the N1 (90- 130ms at FCz), of the early-P3 (220-260ms at Cz) and of the late-P3 (320-360ms at Fz and CPz) following a distractor was fitted to a linear model with Gaussian family distribution model with three within-subject fixed-factors: Music (2 levels: highA music, lowA music), Cue (2 levels: informative, uninformative), DisPosition2 (2 levels: early, late) and DisContent (2 levels: highA dis, lowA dis).

For statistical analysis of target-related potentials (no dis trials), the mean amplitude per subject of the N1 (110-150 ms at FCz) and of the target-P3 (250-500 ms at CPz) following a target was fitted to a linear model with Gaussian family distribution model with two within-subject fixed-factors: Music (2 levels: highA music, lowA music) and Cue (2 levels: informative, uninformative).

## Declarations

### Ethics approval statement

The study was approved by a national ethical committee, Comité de Protection des Personnes Ile de France V, under the number: 2021-A02248-83. All participants gave written informed consent (according to the Declaration of Helsinki).

### Availability of data and materials

The dataset supporting the conclusions of this article will be available in the OSF repository of the corresponding author.

## Acknowledgements

We thank Rémy Masson and Victor Collombat for support in piloting the experiment and Benjamin Morillon for advice on the framing of the manuscript.

