## Supplementary Material for "The dynamic interplay between tonic and phasic arousal shapes attention to optimize performance"

Supplementary Material & Methods

Sound selection and processing

Music excerpts

The 30s music excerpts (50ms rise-time and 50ms fall-time) were extracted from pieces of classical music categorized and rated in Trost et al. (2012) according to their arousing and emotional properties. Only excerpts categorized as emotionally positive were chosen (valence superior to 6, from a 10-point scale with 0 = low pleasantness and 9 = high pleasantness). Twelve low arousing music excerpts (low arousing music: *lowA music*) were selected from the tenderness and peacefulness categories showing an arousal content inferior to 3, from a 10-point scale with 0 = very calming and 9 = very arousing. Twelve highly arousing music excerpts (high arousing music: *highA music*) were selected from the wonder, joy and power categories showing an arousal content superior to 6, from the 10-point scale. When necessary, each excerpt was re-sampled at 44100Hz on 16 bits, edited to last 30 s (50ms rise-time and 50ms fall-time), and converted from stereo to mono with Audacity (GNU GPL v2). Then, the volume was RMS (for “root mean square”) normalized with MATLAB (The MathWorks Inc., 2022). To amplify the contrast between music conditions, the low and high arousing excerpts were binaurally presented in earphones at 40 and 45dB SL, respectively. The two categories of music were compared for tempo, tempo standard deviation (tempo SD), pulse clarity, brightness and roughness of the sound using the MIR toolbox in MATLAB (Lartillot & Toiviainen, 2007), mean values and BF_10_ obtained from Bayesian independent samples t-tests for lowA vs highA music are presented in Sup. Table 1.

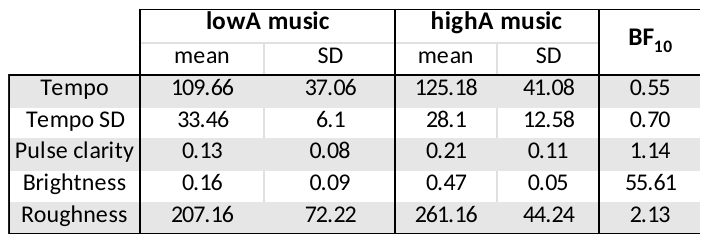

**Sup. Table 1.** Characteristics of the low arousing (lowA) and high arousing (highA) music excerpts. SD: Standard deviation. BF: Bayes Factor.

Distracting sounds

The distracting sounds were chosen from a pool of sounds from Bidet-Caulet et al. (2015) and Widmann et al. ( 2018). When necessary, these sounds were re-sampled at 44100Hz on 16 bits and edited to last 300ms with 5ms rise-time and 20ms fall-time, with Adobe Audition Software (Adobe) or Audacity (GNU GPL v2). In order to modulate phasic arousal, the distracting sounds were previously rated by 26 participants on a 9-point scale with the Self-Assessment Manikins (Bradley & Lang, 1994) for their arousal (calm to exciting), valence (unhappy to happy) and familiarity (non-identifiable to identifiable) contents. Based on these ratings, ten sounds from Bidet-Caulet et al. (2015), and fourteen from Widmann, et al. (2018) , were selected to maximize the difference in arousal content while minimizing the difference in familiarity. It was not possible to minimize the difference in emotional valence since valence and arousal are quite correlated. The twelve low arousing and emotionally positive distracting sounds (*lowA dis*) were rated on average 5.76 ± 0.47, 5.93 ± 0.35, 5.41 ± 0.92, and the 12 high arousing and emotionally negative distracting sounds (*highA dis*) were rated 7.77 ± 0.33, 3.69 ± 0.4, 5.25 ± 1.37 for arousal, valence, and familiarity, respectively.

Screening

Before the first experimental session, participants filled out a short questionnaire about their socio-demographic characteristics. They also completed the adults ADHD Self-Report Scale (Kessler et al., 2005) assessing attention difficulties where the mean score (27.37 ± 7.9) confirmed our population did not present attentional difficulties.

In addition, a sleep questionnaire was fulfilled at the beginning of each experimental session to assess the sleep quality on the night before experimental session. All included participants reported having slept satisfactorily the night before the experiment.

At the end of each block, a drowsiness rating scale (Maldonado et al., 2004) was displayed on the screen: five cartoon faces representing different drowsiness levels were presented, and the participant was asked to choose the one describing best his/her current state. Each cartoon face was then associated with a number from 0 (awake) to 3.60 (sleepy) according to Maldonado et al. (2004) recommendations. Participants ratings indicated that they were moderately drowsy (mean = 1.43, SEM = 0.05).

Supplementary Figures

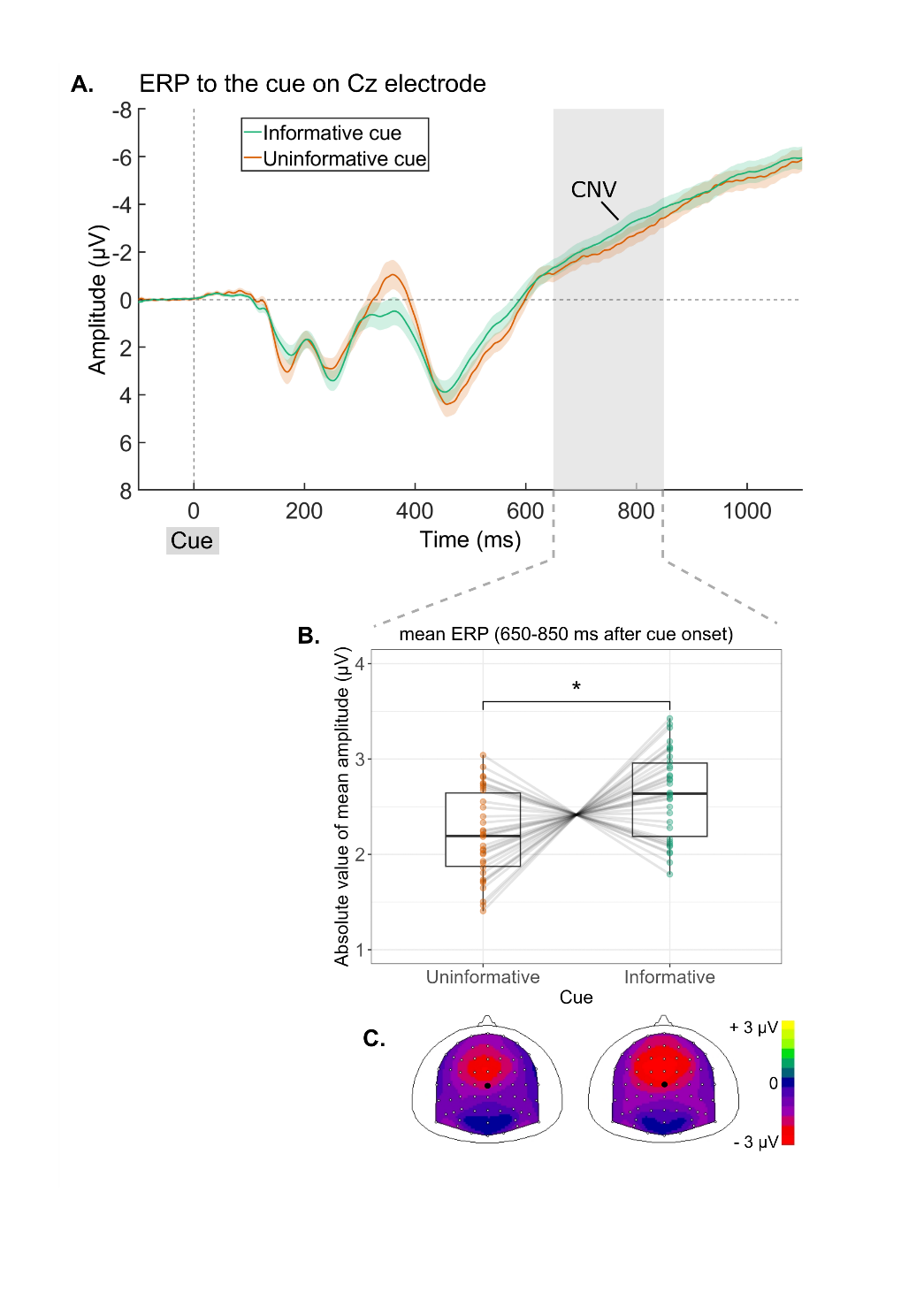

**Sup. Fig. 1.** **Event-Related Potentials (ERP) to the visual cue**, as a function of cue type. **A.** ERP time-course at Cz electrode. Shadowed areas represent standard errors of the mean. ERP time course. Grey areas correspond to the time-window where the main effect of cue was significant. **B*.*** Mean CNV amplitude at Cz in the 650 – 850 ms time window post cue. **C.** Corresponding scalp topographies (top view). The black dot indicates the Cz electrode. Boxplots were plotted using normalized individual data (raw data - subject mean + group mean). Within each boxplot, the horizontal line represents the group median, the lower and upper hinges correspond to the first and third quartiles. The upper whisker extends from the hinge to the largest value no further than 1.5 * IQR from the hinge (IQR = inter-quartile range, or distance between the first and third quartiles). The lower whisker extends from the hinge to the smallest value at most 1.5 * IQR of the hinge. Superimposed to each boxplot, the colored dots represent individual means. * p < 0.05.

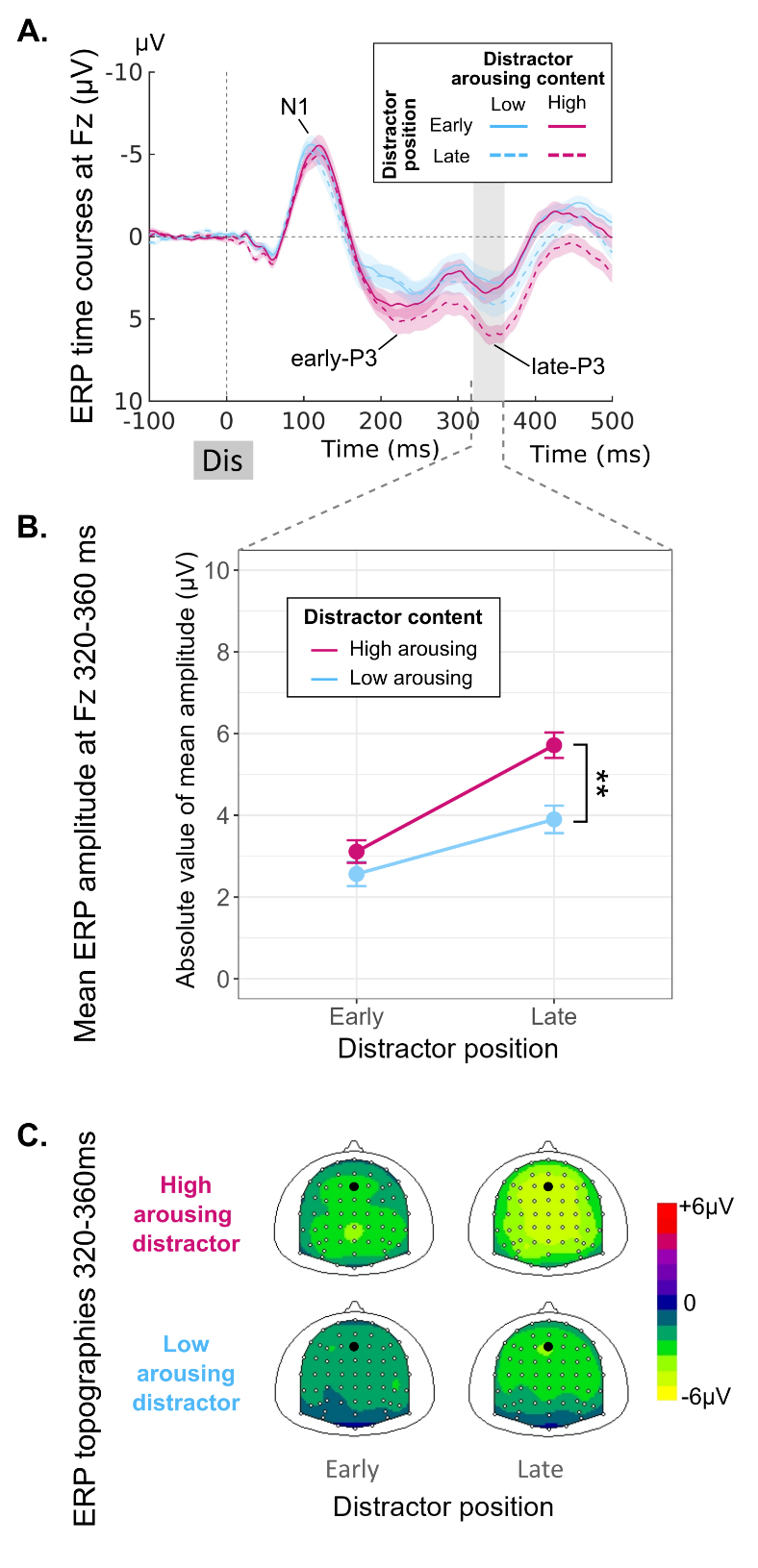

**Sup. Fig. 2.** Event-Related Potentials (ERPs) on Fz electrode to the presentation of a distractor as a function of distractor position (early vs late) and content (low arousing dis vs. high arousing dis). **A.** ERP time-course at Fz electrode. Shadowed areas represent standard errors of the mean. Grey areas correspond to the late-P3 time-window. **B.** ERP mean values for the late-P3 at Fz in the 320 – 360 ms time-window post distractor. **C.** Corresponding scalp topographies (top view). The larger black dot indicates the Fz electrode.

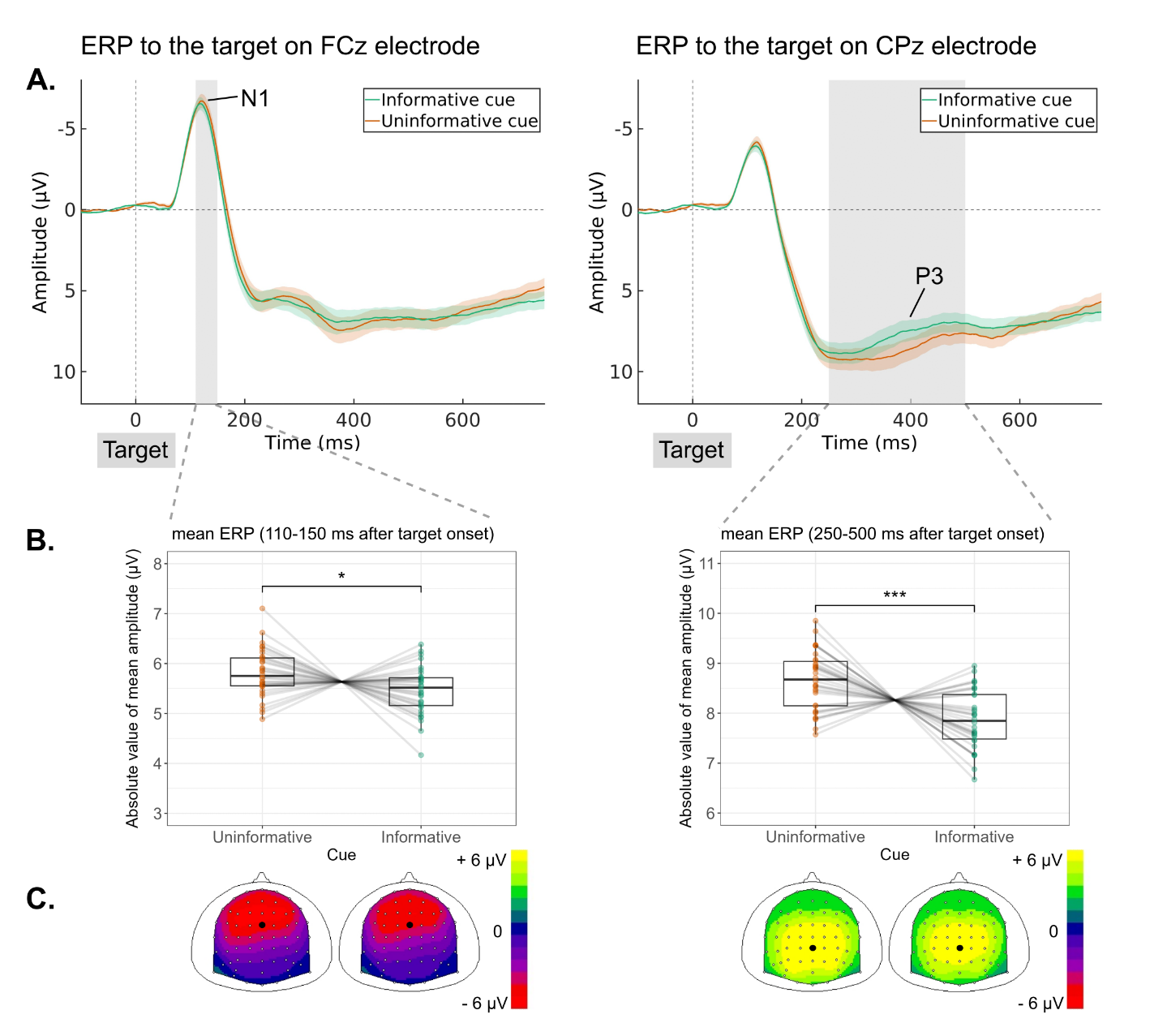

**Sup. Fig. 3.** **Event-Related Potentials (ERP) to the target sound**, as a function of cue type. **A.** ERP time course at FCz and CPz electrodes. Shadowed areas represent standard errors of the mean. Grey areas correspond to the time-window where the main effect of cue was significant. **B.** ERP mean values for the N1 at FCz in the 110 – 150 ms (left), and for the target-P3 at CPz in the 250 – 500 ms (right) time window post target. **C.** Corresponding scalp topographies (top view). The black dot indicates the FCz or CPz electrode. Boxplots were plotted using normalized individual data (raw data - subject mean + group mean). Within each boxplot, the horizontal line represents the group median, the lower and upper hinges correspond to the first and third quartiles. The upper whisker extends from the hinge to the largest value no further than 1.5 * IQR from the hinge (IQR = inter-quartile range, or distance between the first and third quartiles). The lower whisker extends from the hinge to the smallest value at most 1.5 * IQR of the hinge. Superimposed to each boxplot, the colored dots represent individual means. * p < 0.05.
